# Chlamydia trachomatis deploys sphingolipids for genome organisation

**DOI:** 10.64898/2026.04.29.721357

**Authors:** Marcel Rühling, Fabienne Wagner, Till Epprecht, Paul F. Köhling, Fabian Schumacher, Stefan Sachs, Louise Kersting, Julian Fink, Lina Girndt, Burkhard Kleuser, Markus Sauer, Jürgen Seibel, Gregor L. Weiss, Thomas Rudel

## Abstract

*Chlamydia trachomatis* is an obligate intracellular bacterial pathogen and a leading cause of sexually transmitted infections worldwide. During its biphasic developmental cycle, infectious, non-replicative elementary bodies alternate with replicative reticulate bodies within a membrane-bound intracellular niche known as the inclusion. *C. trachomatis* relies heavily on host-derived metabolites, including sphingolipids, which are essential for inclusion integrity, bacterial growth and production of infectious progeny. Here, using expansion microscopy, we uncover an unexpected localization of sphingolipid derivatives within the highly condensed DNA nucleoids of elementary bodies. These sphingolipids are released from nucleoids prior to DNA decondensation during the elementary-to-reticulate body transition, the earliest phenotypic event in the complex developmental cycle of these bacteria. Thereafter, nucleoids undergo a characteristic DNA decondensation process that we visualized by expansion microscopy. By combining super-resolution imaging with a FRET-based metabolic tracking approach and lipidomics, we identified sphingomyelin derived from the sphingolipid analogues as the sphingolipid species predominantly associated with the condensing nucleoids of elementary bodies. Notably, reticulate bodies arrested in their developmental stage fail to accumulate sphingomyelin, suggesting a role for this lipid in stage-specific DNA condensation. Together, our findings suggest an unanticipated role for sphingolipids in bacterial DNA organization and developmental regulation in *C. trachomatis*.

## Introduction

Sphingolipids are essential components of eukaryotic membranes and also function as bioactive signalling molecules.^1^ They regulate diverse cellular processes for instance by acting as ligands for receptors, such as sphingosine-1-phosphate^2^, or as components of lipid microdomains (also called lipid rafts) that influence protein distribution at the host cell surface.^3–7^ Dysregulation of cellular sphingolipid levels are associated with diseases^8, 9^ underscoring the importance of maintaining physiological sphingolipid levels through a tightly regulated metabolic network.^1^ Central to sphingomyelin (SM) catabolism are sphingomyelinases (SMases), which hydrolyse SM by removing its polar phosphocholine head group to generate ceramide.^10^ In humans, SMases are classified according to their pH optimum as neutral^11^, acidic^12, 13^ or alkaline.^14^ These enzymes play essential roles in regulating membrane composition and signal transduction^15, 16^ and have also been implicated in promoting bacterial and viral infections.^17–21^ SMases constitute important virulence factors in several bacterial pathogens, including *Staphylococcus aureus*^22^ and *Chlamydia pneumoniae.*^23^

The obligate intracellular bacterium *Chlamydia trachomatis* is a major cause of sexually transmitted diseases.^24^ *C. trachomatis* exhibits a biphasic developmental cycle comprising infectious elementary bodies (EBs) and metabolically active, replicative reticulate bodies (RBs).^25, 26^ Upon adherence to host cells, EBs interact with host receptors and inject effector proteins into the host cell cytosol via the type three secretion system (T3SS), thereby triggering actin polymerization and bacterial internalization into a membrane-bound vacuole.^27–30^ Once inside the host cell, EBs differentiate into RBs and establish a large replicative compartment, the so called inclusion. At later stages of the intracellular life cycle, RBs redifferentiate into EBs, which are ultimately released from host cells to disseminate the infection.^25, 26^ During RB-EB redifferentiation, histone-like proteins mediate DNA condensation and formation of a chlamydial nucleoid, a process that is essential for transcriptionally down regulating gene expression in the metabolically less active EB state. Conversely, nucleoid disassembly during EB-to-RB differentiation initiates gene expression.^26, 31–33^ While recent efforts provide mechanistic insights into RB-to-EB transformation^34^, the process of how EBs develop into RBs is mostly unknown.^25^ Currently, it is thought that EB-to-RB and RB-to-EB differentiation occurs via poorly defined intermediate developmental stages, the so-called intermediate bodies.^25, 34–36^

The chlamydial genome only encodes ∼1000 genes^37^ and thus lacks important metabolic pathways. Therefore, *C. trachomatis* is highly dependent on host cell metabolites including amino acids^35^, nucleotides^38^ and lipids.^39^ Sphingolipids have been shown to be particularly important in *C. trachomatis* infection.^40–43^ For instance, they are acquired by the bacteria via the ceramide transfer protein (CERT) that is recruited to the chlamydial inclusion. Subsequently, sphingolipids are incorporated into bacterial and inclusion membranes.^39, 42–46^ SM, the most abundant sphingolipid in human cells, was demonstrated to stabilize chlamydial inclusions and facilitates replication.^40, 41^ SM can be acquired by redirecting exocytic vesicles originating from the Golgi apparatus, the site of SM *de novo* synthesis, to the inclusion membrane.^47^ Moreover, SM formation was observed in absence of the host SM biosynthesis machinery during *C. trachomatis* infection, suggesting the existence of a chlamydial SM synthase and highlighting the importance of this particular sphingolipid species for the bacteria.^48, 49^ By contrast, other sphingolipids, such as sphingosine, possess bactericidal properties and play a role for the host immune response to *C. trachomatis* and other pathogens.^50^ ^46, 51^

The diffraction limit of light restricts the spatial resolution of conventional light microscopy to ∼250 nm.^52^ Introduced in 2015, expansion microscopy (ExM) is a super-resolution imaging technique that can circumvent this fundamental limitation.^53^ During ExM, a fixed specimen is embedded into a swellable hydrogel that, upon contact to water, expands and thereby the spatial distance between individual fluorophore is mechanically enlarged resulting in an enhanced resolution achievable on conventional microscopes. The so-called expansion factor is defined by the gel composition and defines how much a gel can expand in each spatial direction (e.g. 4-fold or 8-fold, referred to as 4xExM or 8xExM, respectively).^54, 55^ Proteins can be efficiently anchored into the gel matrix and can be visualized by antibodies^56^ or fluorescently tagged N-hydroxysuccinimide esters (NHS esters), in so-called pan-ExM.^57^ By contrast, most naturally occurring lipids are not embedded into the hydrogel and the limited option to stain lipids poses additional challenges. However, we^44, 45^ and others^58^ developed protocols that enable ExM of functionalized lipid derivatives.

Here, we used a variety of imaging modalities to investigate SM distribution during the chlamydial life cycle. Unexpectedly, we observed highly dynamic incorporation and removal of an SM derivative within bacterial nucleoids during the developmental cycle, suggesting a previously unrecognized role for SM in DNA organization.

## Results

Sphingolipids are essential for replication and the formation of infectious progeny during *C. trachomatis* infection; ^40, 41^ however, their specific function within *Chlamydia* remains unknown. We previously introduced two trifunctional sphingomyelin derivatives (TFSM1 and TFSM2) that enable visualization of SM distribution at nanoscale resolution via 4x ExM (**Supp. Fig. 1**, **Supp. Note 1**).^45, 46^ We here optimized these tools to further investigate the distribution and metabolization of TFSMs throughout the entire chlamydial developmental cycle.

### EBs possess a cell pole enriched with TFSM

Visualizing TFSMs by 4x ExM allowed detection of individual bacteria within an inclusion, while visualization of structures within EBs is challenging due to their small size of 200-300 nm. To investigate the lipid distribution within EBs, we combined TFSM2 labeling with 8x ExM^55, 59^ thereby further enhancing the spatial resolution achievable on conventional microscopes. This approach revealed a characteristic lipid accumulation at a single site within EBs (**Fig. 1**, a, b). We excluded the possibility that this pattern is caused by staining or expansion artifacts by analyzing expansion factors and signal-to-background ratios (**Supp. Fig. 2**, **Supp. Note 2**).

**Fig. 1.**
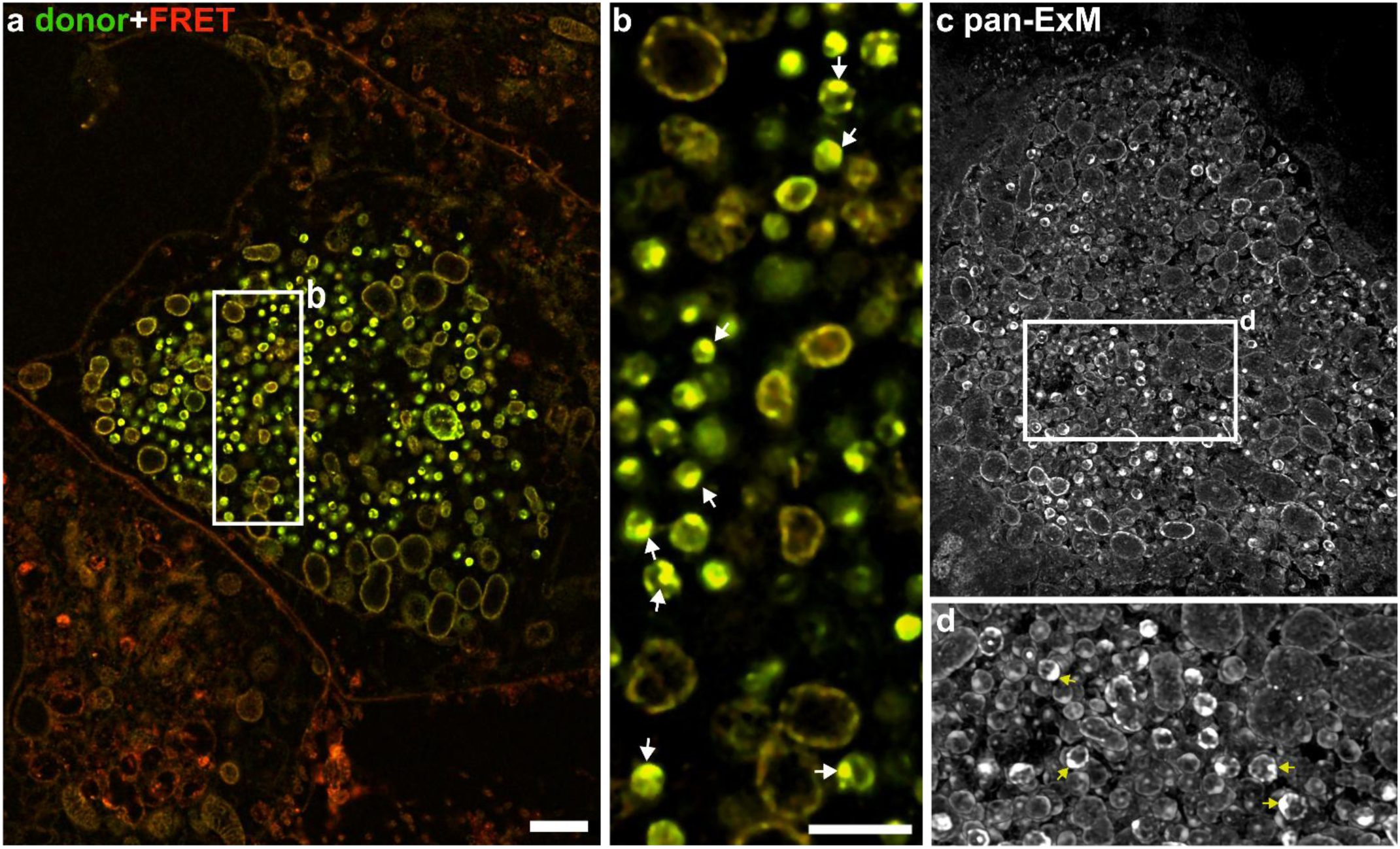
TFSM accumulates at a specific site within EBs. HeLa 229 cells were infected with *C. trachomatis* in presence (a, b) or absence (c, d) of TFSM2 for 32 h and stained for TFSM2 (a, b) or proteins by an NHS ester (c, d). Samples were visualized by 8x ExM. c and d show a 3D reconstruction. Zoomed area is indicated by a white rectangle in a and c. TFSM2 (b) or protein (d) accumulation sites are marked by arrows. Scale bar: 8 µm (∼1 µm with 8-fold expansion factor).

In certain cases, ExM can cause deformation of bacterial particles, when protocols are not carefully adjusted.^60^Additionally, the treatment of infected cells with TFSMs could cause the accumulation. To excluded these potential sources for artifacts, we combined 8x ExM with pan-ExM, an approach based on NHS ester staining of endogenous proteins.^57^ This strategy enables the visualization of the sample ultrastructure independently of TFSM labeling (**Fig. 1**, c). By pan-ExM, we did not only demonstrate the structural integrity of individual chlamydial particles but also detected a similarly localized accumulation to that observed in TFSM2-labeled samples (**Fig. 1**, d), indicating that one pole of the EB is enriched in both sphingolipids and proteins. This polar enrichment was confirmed by 3D reconstructions of individual EBs (**Fig. 2**, a).

**Fig. 2.**
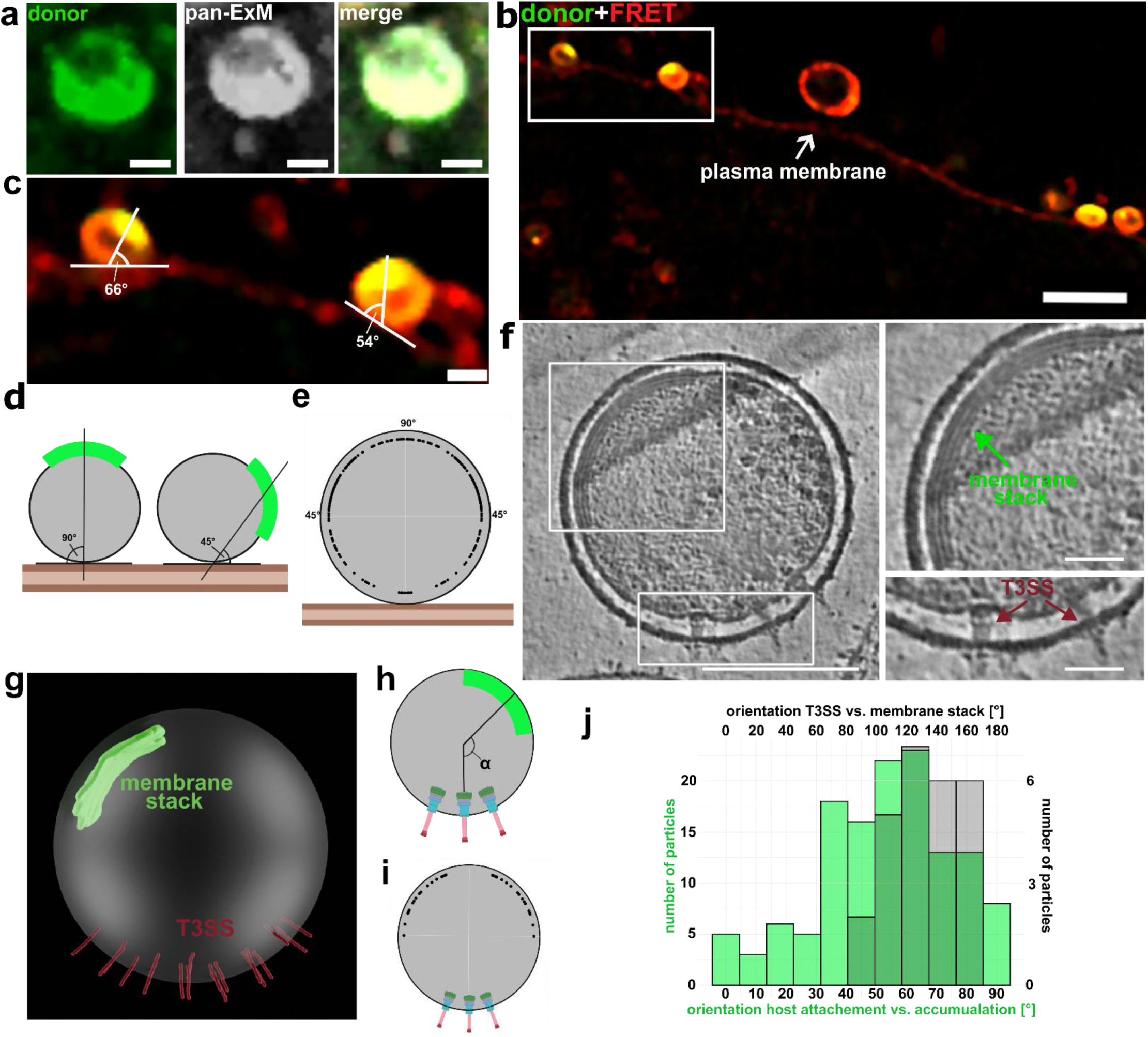
EBs possess a TFSM-enriched cell pole opposed to the T3SS. **a, TFSM is enriched at one EB hemisphere.** 3D reconstruction of a single EB stained for TFSM and whole protein (pan-ExM) recorded by 8x ExM. Scale bar: 1 µm (∼125 nm with 8x expansion factor). **b-e, lipid-enriched membrane sites are located distal from the plasma membrane during EB adherence.** HeLa 229 cells were infected with *C. trachomatis* in the presence of TFSM2 for 1 h to allow EBs to adhere to host cells. Cells were washed and imaged by 8x ExM (b, c). Tangent on the EB-plasma membrane contact site was delineated and the angle between the tangent and the accumulation was determined (c, d). Individual angle measurements are visualized as dots in e. Scale bar: 5 µm (b, 625 nm with 8x expansion factor) and 1 µm (c, 125 nm with 8x expansion factor). **f-i, T3SS and membrane stacks are located at opposing sites of EBs.** Cryo-electron tomogram of an whole cell EB (f) and 3D segmentation used for analysis (g). Angles between T3SS and the membrane stack were determined in the 3D segmentation (h). Individual angle measurements between T3SSs and membrane stack are visualized as dots in i. *n*=26 EBs. Scale bar: 200 nm (whole EB) and 50 nm (magnified images). **j, membrane stacks are enriched in sphingolipids.** The orientations between the T3SS and membranes stacks (cryo-ET, grey; f-i) and between the EB-plasma membrane contact site and the lipid accumulation (8x ExM, green; b-e) is compared.

The asymmetric accumulation of lipids to one pole prompted us to analyze the orientation of EBs during attachment to the host plasma membrane by 8x ExM (**Fig. 2**, b-e, **Supp. Movie 1**). The lipid accumulation was typically observed at the EB pole opposing the host cell during attachment. To quantify its spatial orientation, we defined the tangent at the point of EB-plasma membrane attachment and determined the angle between this tangent and the lipid accumulation (**Fig. 2**, c and d). The analysis demonstrated a strong bias toward localization distal to the plasma membrane, with the TFSM-enriched hemisphere of the EB exhibiting an average orientation angle of ∼60° (**Fig. 2**, e).

To further elucidate the EB architecture, we visualized whole-cell and cryo-focused ion beam (FIB) thinned EB preparations by cryo-electron tomography (cryo-ET, **Fig. 2**, f, g). This analysis revealed that the T3SS is predominantly localized to one bacterial cell pole (n = 105 EBs), while characteristic membrane stacks were positioned at the opposing pole in 71 % of the analyzed cells (**Fig. 2**, f, g, **Supp. Movie 2**) consistent with previous observations.^27^ The absence of the membrane stack in the remaining tomograms likely reflects technical limitations of cryo-EM, such as cell thinning during sample preparation or missing-wedge artifacts, rather than true biological heterogeneity. Importantly, EB preparations with and without TFSM treatment exhibited indistinguishable ultrastructural features, indicating that TFSM labeling does not measurably alter EB architecture under the conditions used in this study.

We quantified the orientation of the membrane stack in relation to the T3SS by determining the angle between the two structures (see, **Fig. 2**, h), which yielded an average of ∼135° (**Fig. 2**, i). We assume that the T3SS is located in close proximity to the plasma membrane during adherence to host cells.^27^ By comparing the orientation between the T3SS and the membrane stack (cryo-ET), and the orientation between EB-host attachment sites and the TFSM-enriched cell pole (8x ExM), we concluded that the TFSM-enriched EB pole also contains additional membrane stacks (**Fig. 2**, j). Taken together, our data suggest that EBs possess a polarized architecture in which one cell pole is enriched in sphingolipid derivatives and associated membrane structures, while the opposing cell pole contains the T3SS that contacts host cells during attachment. The spatial segregation of these features implies that the sphingolipid-enriched pole is unlikely to play a direct role in host cell adhesion and bacterial uptake. Instead, it may serve a distinct function during the chlamydial developmental cycle, which we sought to investigate in the present study.

### Sphingolipid derivatives are incorporated into chlamydial nucleoids

The membrane stacks are usually aligned with the inner EB membrane, which explains a general increase in lipids detected at this cell pole (**Fig. 2**, f, g). However, in our 8x ExM samples, we frequently observe a globe-shape TFSM-enriched structure that protruded into the cytosol of the EB (**Fig. 1**, a; **Fig. 3**, d, e). This morphology could not be readily explained by membrane stacks alone. Comparison of 8x ExM and cryo-ET images revealed a striking correspondence between these structures. The region occupied by the TFSM-positive protrusion in 8x ExM images coincided with a rather homogenous electron-dense and ribosome-depleted area in cryo-electron tomograms of cryo-FIB thinned EBs. Previous studies have identified this region as the chlamydial nucleoid^27^, which contains highly condensed bacterial DNA (**Fig. 3**, a, b, **Supp. Movie 2**). These observations therefore suggest that sphingolipid derivatives are associated not only with the membrane-rich pole of the EB but also with the nucleoid region.

**Fig. 3.**
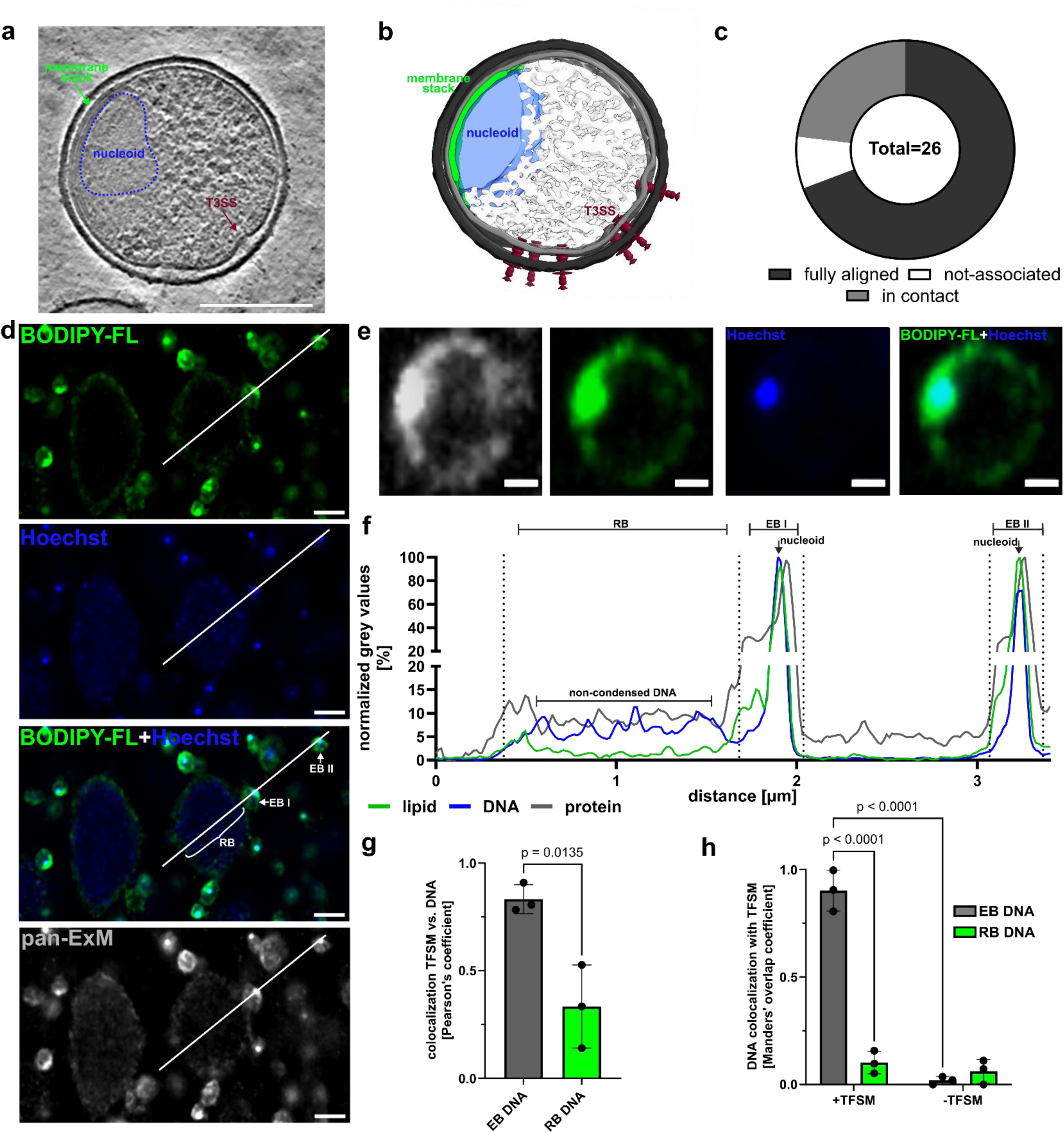
TFSM is incorporated into chlamydial nucleoids. **a-c, Cryo-ET reveals a close association of the chlamydial nucleoid with membrane stacks.** An EB preparation was plunge-frozen, thinned by cryo-FIB milling, imaged by cryo-ET and a representative tomogram (a) was 3D-segmented (b). Scale bar: 200 nm. Association of nucleoids with additional membrane stacks in tomograms was categorized as “fully aligned”, “in contact” and “not-associated” (c). **d-f, TFSM2 colocalizes with EB DNA.** HeLa 229 cells were infected with *C. trachomati*s in the presence of TFSM2 and visualized by 8x ExM. TFSM2 was stained by BODIPY-FL-DBCO. DNA and proteins were stained by Hoechst and a fluorescent NHS ester, respectively (d, e). A plot profile of a selected area (white line in d) was measured, and particles/nucleoids were marked by arrows and brackets (f). Scale bar: 4 µm (d) and 0.8 µm (f), (0.5 µm and 0.1 µm with 8-fold expansion factor, respectively). **g, h, TFSM2 colocalizes with EB DNA.** HeLa 229 cells were infected with *C. trachomati*s in the presence or absence of TFSM2 for 32 h and imaged by 8x ExM. Regions containing EBs or RBs were selected and analyzed independently for Pearson’s (g) or Manders’ overlap coefficient (h). Statistics: Unpaired Student’s t-test (g) or two-way ANOVA and uncorrected Fisher’s LSD (h).

In 69 % of the analyzed EBs in these tomograms, in which we detected both the nucleoid and the membrane stack (*n* = 26), the nucleoid was fully aligned to the membrane stack, whereas in an additional 23% of the cells the nucleoid was slightly offset but still remained in contact with the membranes (**Fig. 3**, b, c). In two cells (8%) no apparent association between nucleoid and membrane stack was detected. Thus, we hypothesized that the TFSM accumulation co-localizes with the chlamydial nucleoid.

To determine the spatial relationship between EB nucleoids and the TFSM accumulation, we performed 8x ExM and stained TFSM2 with BODIPY-FL-DBCO, proteins by pan-ExM and DNA with Hoechst. To assess the degree of spatial overlap, we generated plot profiles of different chlamydial particles. This analysis revealed a striking colocalization of TFSM2 and DNA (**Fig. 3**, d, e). Interestingly, this colocalization was specific to EB nucleoids and was not observed in RBs, in which DNA appeared non-condensed and distributed throughout the bacterial cell (**Fig. 3**, d, f).

Because click-labeling dyes can produce substantial background fluorescence, we assessed whether the observed nucleoid signal could result from staining artifacts. Tho this end, we measured the mean BODIPY-FL fluorescence in individual nucleoids in samples infected in presence or absence of TFSM2 (**Supp. Fig. 3**, a, b, **Supp. Note 3**). Nucleoids in TFSM2-treated samples displayed significantly higher fluorescence, excluding staining artifacts originating from the click dye. Colocalization of DNA with α-NH_2_-ω-N_3_-C_6_-ceramide, a molecule developed for membrane visualization by ExM,^44^ further confirmed the close association of lipids with the chlamydial nucleoids (**Supp. Fig. 3**, b. **Supp. Note 3**). To quantify the TFSM-DNA colocalization, we determined Pearson’s (**Fig. 3**, g) and Manders’ (**Fig. 3**, h) coefficients between Hoechst and BODIPY-FL channels in regions containing either EBs or RBs. Both measurements showed a strong correlation between the lipid and EB nucleoids but were markedly lower for non-condensed RB DNA. Strikingly, ∼90% of nucleoid areas also contained TFSM, an effect observed only in the presence of TFSM, again excluding staining artifacts of the click dye. These results provide quantitative support for the co-localization of the TFSM accumulation with condensed chlamydial DNA. Based on these observations, we hypothesized that TFSM-derived lipids accumulate within, or in immediate proximity to, the chlamydial nucleoid.

As TFSMs are designed sphingolipid derivatives, we next visualize endogenous membranes in relation to the chlamydial nucleoid. Therefore, we tested whether the recently developed ExM-compatible membrane probe pGk13a^61^ associates with the nucleoid. pGk13a enabled visualization of cellular membranes and individual bacterial particles via 8x ExM (**Supp. Fig. 4**, a and b). Similar to TFSM, we detected a pGk13a accumulation on one EB pole that closely associates with the nucleoid (**Supp. Fig. 4,** c-e) and supposably corresponds to the membrane stack detected by cryo-ET. Although, pGk13a and Hoechst signal completely overlapped in several particles (**Supp. Fig. 4,** c and d), we frequently observed that the signals only partially colocalize. Additionally, we did not detect the cytosolic globe-shaped lipid protrusion, which was characteristic for samples visualized by TFSM.

These data support our hypothesis that membrane stacks build an interphase between the inner membrane and the nucleoid. At the same time, the more extensive distribution of TFSM-derived signals suggest that sphingolipid derivatives may occupy regions beyond those visualized by pGk13a.

### Lipid derivatives are released from nucleoids early during EB-to-RB differentiation

Next, we elucidated the association of TFSM within the bacterial nucleoids in the early developmental cycle of *C. trachomatis*. Upon host cell entry, EBs differentiate into RBs, a process that is poorly understood.^25^ Because DNA in RBs is decondensed, nucleoids must be dissembled during EB-to-RB differentiation. To investigate the dynamics of this transition, we infected HeLa 229 cells preloaded with TFSM2 and visualized lipid and DNA during the first 6 h of infection by 8x ExM (**Fig. 4**, a). Within the first hour p.i., most bacterial particles were either attached to the host plasma membrane or located at the cell periphery, indicating that invasion had occurred recently (see also **Fig. 2**, b, c). By 3 h p.i., bacteria clustered in small groups, and by 5 h p.i. most bacteria were located at one single intracellular site adjacent to the host nucleus. These observations are consistent with previous reports showing the localization of chlamydial inclusions at the peri-Golgi and microtubule organization center (MTOC).^47, 62, 63^

**Fig. 4.**
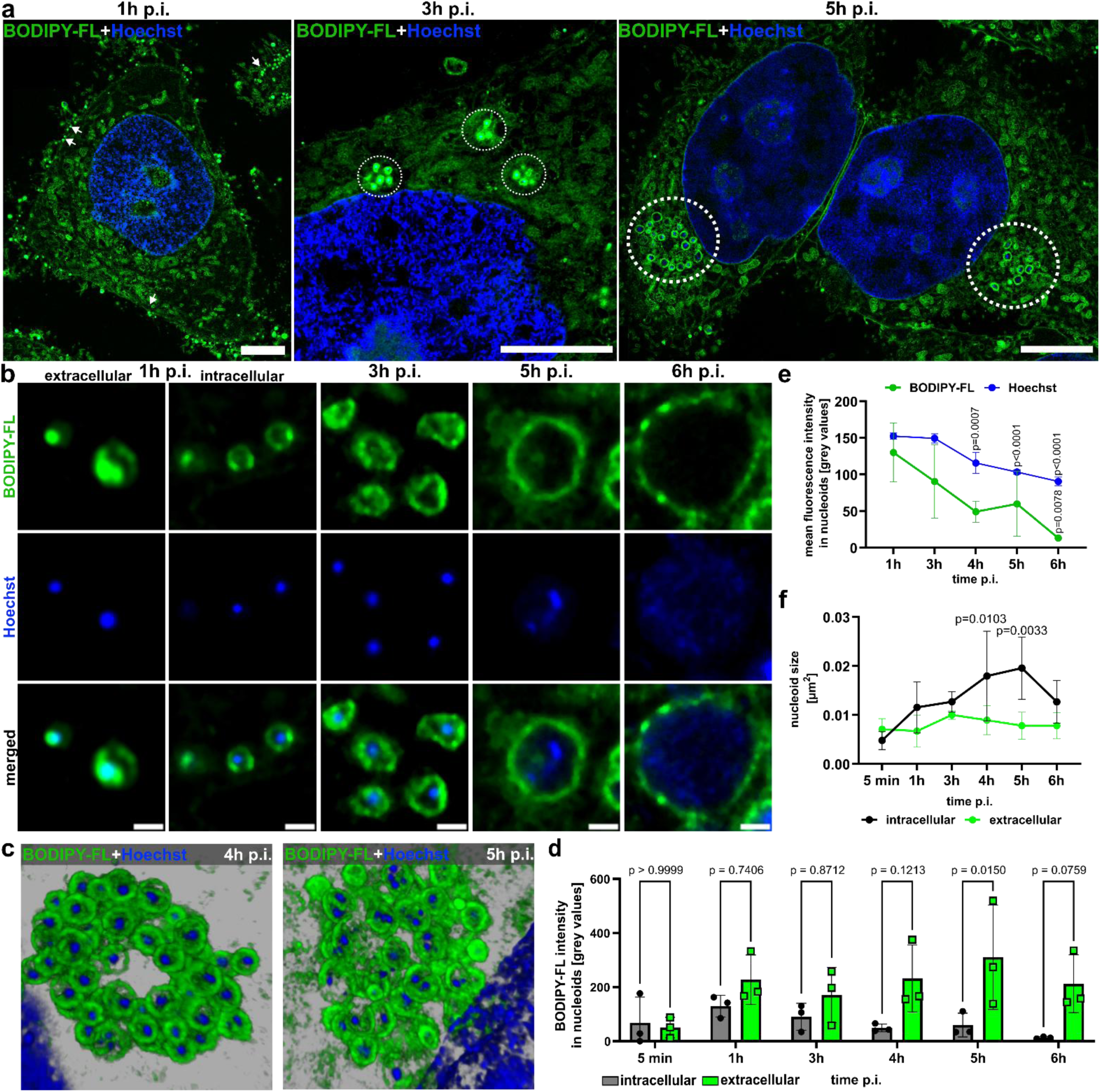
Lipid-DNA association is resolved during EB-RB differentiation. **a, Chlamydial particles gather at a certain site in host cells.** HeLa 229 cells were infected with *C. trachomatis* in the presence of TFSM2 and fixed at the indicated time points. The samples were stained with BODIPY-FL-DBCO and 8-fold expanded. DNA was visualized with Hoechst. Particles are marked with white arrows (1 h p.i.) or circles (3 and 5 h p.i.). Scale bar: 40 µm (∼5 µm with 8-fold expansion factor). **b, c Visualization of EB-to-RB differentiation during the first 6 h of infection**, Samples were prepared as described in a and fixed at the indicated time points. Images were reconstructed in 3D (c). Scale bar: 2 µm (∼250 nm with 8-fold expansion factor). **d-f, Lipid contend of nucleoids decreases during DNA decondensation,** regions containing nucleoids in intracellular or extracellular bacteria were selected in Hoechst channels. Mean intensities of BODIPY-FL (d, e) and Hoechst (e) in selected nucleoids as well as the size (f) of the selected nucleoids were measured. P-values refer to comparison between the indicated time point and 1h (e) or 5 min (f). n=3. Statistics: Two-way ANOVA and Šídák’s (d) or Tukey’s multiple comparison (e), one-way ANOVA and Dunnett’s multiple comparison (f).

We next analyzed individual bacteria over time (**Fig. 4**, b). At 1 h p.i., extracellular EBs attached to the plasma membrane retained TFSM2 within nucleoids, which remained associated with the inner bacterial membrane. In contrast, intracellular bacteria frequently exhibited morphological changes, with condensed nucleoids devoid of TFSM and relocated to the center of the bacterial particle. These observations indicate that dissociation of the lipid-DNA association and detachment of the nucleoid from the bacterial envelope are among the earliest detectable steps in EB-to-RB differentiation. By 3 h to 4 h p.i., the bacterial DNA still appeared at a single location, but subsequently adopted a characteristic decondensation pattern, frequently appearing as multiple distinct foci (**Fig. 4**, c). By 6 h p.i., DNA was often completely decondensed and distributed throughout the bacterial cytosol (**Fig. 4**, b), resembling the nucleoid organization of RBs at mid-to-late stages of infection (**Fig. 3**, d). During the first 6 h of infection, particle diameters increased from ∼200 nm to 750 nm, while after 32 h p.i. RBs in average measured 1 µm in size (**Supp. Fig. 3**, c).

To quantify the lipid content of bacterial DNA, we selected individual DNA foci in the Hoechst channel, determined average fluorescence intensities in the lipid (BODIPY-FL) channel and subtracted the background signal detected in samples not incubated with TFSM2 (**Fig. 4**, d, e). Extracellular EBs showed only marginal TFSM2 incorporation after 5 min p.i., while a high TFSM2 signal was detected by 1 h p.i. (**Fig. 4**, d). These results indicate that TFSM2 is taken up by plasma membrane-attached EBs from the host or the environment and subsequently incorporated into the nucleoid. The TFSM content in nucleoids of extracellular bacteria remained constant during the first 6 h of infection, whereas TFSM-DNA association decreased in intracellular bacteria over time (**Fig. 4**, d, e).

To detect DNA decondensation, we measured the Hoechst mean intensity (**Fig. 4**, e) or the size of DNA foci (**Fig. 4**, f) in intracellular bacteria. During the first 5 h of infection, Hoechst intensity decreased while the area of individual DNA foci increased, demonstrating progressive unwrapping of DNA. Significant changes in both measures were detected first at 4 h p.i.. By 6 h p.i., the size of DNA foci had decreased while the total area containing DNA had increased, as DNA became nearly fully decondensed in most particles. These data indicate that nucleoid decondensation during EB-to-RB differentiation is a continuous process that proceeds within the first 6 h of infection through several morphologically distinct stages. These stages can be distinguished by the degree of DNA-TFSM association and the number and distribution of DNA foci.

### Metabolized TFSM enriches in *Chlamydia* inclusions during RB-to-EB redifferentiation

Direct interaction between DNA and zwitterionic lipids, such as SM, has been reporter *in vitro*.^64–71^ However, we previously proposed that the polar phosphocholine head group of TFSMs is primarily cleaved off within *C. trachomatis* inclusions.^45, 46^ This processing should generate the less polar ceramide metabolite, which is less likely to form strong interactions with DNA or proteins and hence, it would be unexpected that ceramide would associated with the nucleoid during maturation of the chlamydial inclusion and formation of infectious progeny. To better understand the nature and fate of sphingolipids during inclusion maturation, we investigated SM metabolism during *C. trachomatis* infection. First, we analysed SM transport to the chlamydial inclusion in general. Therefore, we treated infected HeLa cells with BODIPY-FL-C_12_-SM, a lipid analogue that unlike TFSMs possess a naturally, even though rarely occurring, acyl chain length and hence, should adequately mimic the physiological lipid transport. Direct conjugation of the BODIPY-FL fluorophore to this lipid analogue enables its visualization without further staining procedures, thereby minimizing the risk for dye artefacts.

We observed uptake of this derivative into chlamydial inclusions in both, WT and CERT K.O. cells (**Supp. Fig. 5**, a). This suggests that this SM derivative can reach the inclusion independently of CERT, indicating the presence of vesicular SM transport routes to the *C. trachomatis* inclusion as previously described.^72^ However, imaging of BODIPY-FL-C_12_-SM only demonstrates uptake and localization of the lipid and does not provide information about its metabolization. In contrast, TFSMs can be used not only to visualize SM distribution but also allow monitoring SM turnover with spatial resolution (**Supp. Fig. 1**).^45^ Using click chemistry, TFSMs can be equipped with two fluorophores, one at the headgroup and one at the tail of the lipid analogues. Importantly, the sites where fluorophores are attached to the lipid tail of the molecules differ between TFSM1 (sphingoid backbone) and TFSM2 (acyl side chain). If the chosen fluorophores form a FRET pair, FRET efficiency measurements can be used to detect degradation of TFSMs through SMases by conventional microscopy or 4x ExM.^45^ The FRET efficiency correlates with the concentration of non-metabolized “native” TFSM in the selected area, while a relatively low FRET efficiency indicates predominant presence of a TFSM metabolite. Therefore, TFSMs provide a unique tool to simultaneously monitor sphingolipid localization and metabolism, enabling investigation of the role of sphingolipids during inclusion maturation.

To elucidate the dynamics of TFSM metabolization during infection, we measured the FRET efficiency in inclusions and host cell areas by FRET acceptor bleaching over an infection course of 46 h (**Fig. 5**, a) to determine the native TFSM content of inclusions. We observed similar FRET efficiencies in inclusions and host cell membranes at 16 h p.i., while the FRET efficiency in inclusions decreased at later time points of infection starting at 24 h p.i. (**Fig. 5**, b). TFSM degradation within inclusions was further assessed by normalizing inclusion FRET efficiency to that of the host cell (ratio “inclusion vs. host cell”). Consistently, the FRET efficiency ratio decreased during infection, indicating an enrichment of metabolized TFSM in inclusions compared to the host cell in the mid-late infection cycle. This trend was observed in both unexpanded samples and samples subjected to 4x ExM (**Fig. 5**, c; **Supp. Fig. 5**, b).

**Fig. 5.**
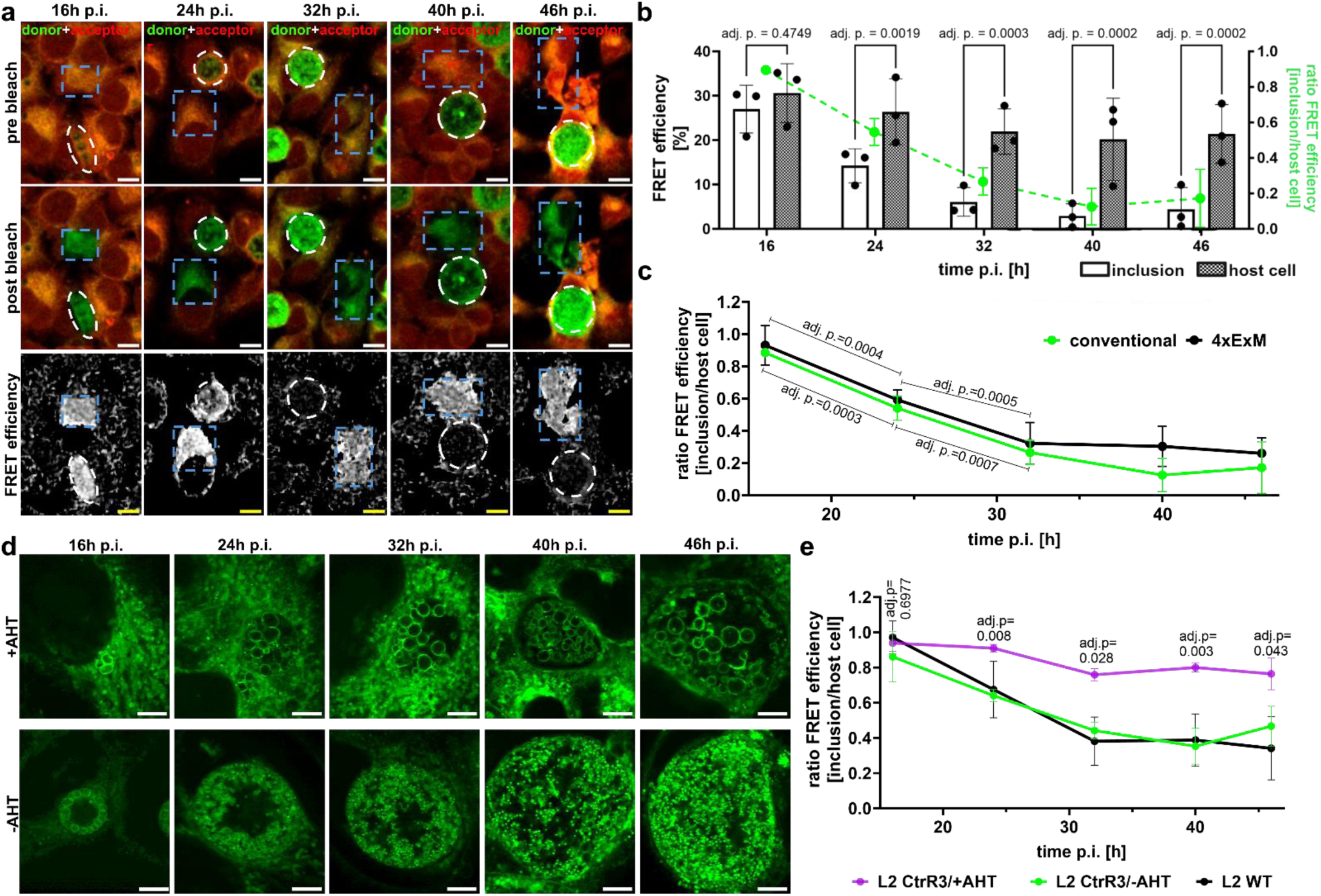
Accumulation of metabolized TFSM in inclusions is associated with RB-EB redifferentiation. **a-c, Metabolized TFSM accumulates in inclusions during maturation.** HeLa cells were infected with *C. trachomatis* for different time points, stained for TFSM2 and analyzed by conventional confocal microscopy (a, b) or 4x ExM (c). FRET acceptor bleaching of host cell membranes (blue rectangles) or chlamydial inclusions (white circles) was performed to determine FRET efficiency. Ratios of FRET efficiencies detected in inclusions vs. host cells were calculated (b, green and c). Scale bars: 10 µm. **d, e, Blockage of RB/EB redifferentiation attenuates accumulation of metabolized TFSM in inclusions.** HeLa cells were infected with *C. trachomatis* serovar L2 CtrR3 and expression of the sRNA CtrR3 was induced by addition of anhydrous tetracycline (AHT) or left uninduced. Samples were stained for TFSM2 and analyzed by 4x ExM (d). FRET acceptor bleaching was performed in regions containing host cell membranes or chlamydial inclusions and ratios “inclusion vs. host cell” were calculated (e) and compared to results of wildtype *C. trachomatis* serovar L2. Scale bars: 20 µm (with 4x expansion factor ∼5 µm). Adj. p values in e relate to comparison of -AHT and +AHT. Statistics: Two-way ANOVA and Šidák’s multiple comparison (b, e) or mixed-effects model (REML) and Tukey’s multiple comparison (c).

Commonly, EB formation initiates at ∼24 h p.i.,^25^ the earliest time point at which we detected accumulation of metabolized TFSM in inclusions. This temporal overlap led us to hypothesize that TFSM is metabolized during RB-to-EB redifferentiation. To identify whether TFSM metabolization is connected to inclusion maturation, we used a *C. trachomatis* strain that conditionally overexpresses the chlamydial sRNA CtrR3, which prevents EB formation.^73^ Anhydrous tetracycline (AHT)-induced expression of CtrR3 during infection of HeLa cells was validated by RT-qPCR (**Supp. Fig. 5**, c) and infection was monitored by 4x ExM (**Fig. 5**, d). In the absence of AHT, inclusions developed comparably to those of the wildtype strain, whereas induction of CtrR3 expression impaired inclusion maturation, most notably by the absence of EBs (**Fig. 5**, d) and the presence of enlarged RBs (**Supp. Fig. 5**, d; **Supp. Note 4**). Next, we measured FRET efficiency in inclusions and host cell areas by acceptor bleaching, which revealed that CtrR3 expression markedly impaired TFSM metabolization (**Fig. 5**, e), evident by similar FRET efficiencies, and thus native TFSM content, in inclusions and host cells. In summary, these data indicate that metabolized TFSM accumulates within chlamydial inclusions during RB-to-EB redifferentiation.

### TFSM is metabolized within chlamydial inclusions

Because *C. trachomatis* strongly depends on host cell metabolites^35, 38, 39^, we hypothesized that TFSM is metabolized by host enzymes and subsequently shuttled into the inclusion. To test this possibility, we investigated whether central host factors of sphingolipid metabolism such as CERT or SMases contribute to TFSM metabolization during *C. trachomatis* infection (**Supp. Fig. 6** and **Supp. Note 5**). We found no evidence for infection-induced upregulation of host SMases, neither at the expression level (**Supp. Fig. 6,** a and b) nor the cellular SMase activity (**Supp. Fig. 6**, c and d). Consistently, we detected, using our FRET-based approach, that metabolized TFSM still accumulated in inclusions upon pharmacological inhibition of neutral SMase (NSM) and/or genetic ablation of acid SMase (ASM, **Supp. Fig. 6**, e). Next, we investigated whether host ceramide transport could be involved in accumulation of metabolized TFSM in inclusions. CERT mediates non-vesicular transport of ceramide and is recruited to the inclusion membrane. Hence, metabolized TFSM could be theoretically channeled into inclusions via CERT. As described previously^43^, infection was impaired in host cells lacking CERT. However, the enrichment of metabolized TFSM within inclusions was still detected in CERT K.O. cells by the FRET-based readout, even though with slightly delayed kinetics.

Since our data did not support a critical role for these host factors in the enrichment of metabolized TFSM, we hypothesized that TFSM is converted within the inclusion. Therefore, we localized metabolized TFSM in individual bacterial particles by imaging inclusions at different time points p.i. by 4x ExM. Overlay images of FRET and donor channels revealed a high proportion of metabolized TFSM in EBs at 24 h and 32 h p.i. as indicated by the green color of these particles compared to RBs and host cells (**Fig. 6**, a).

**Fig. 6.**
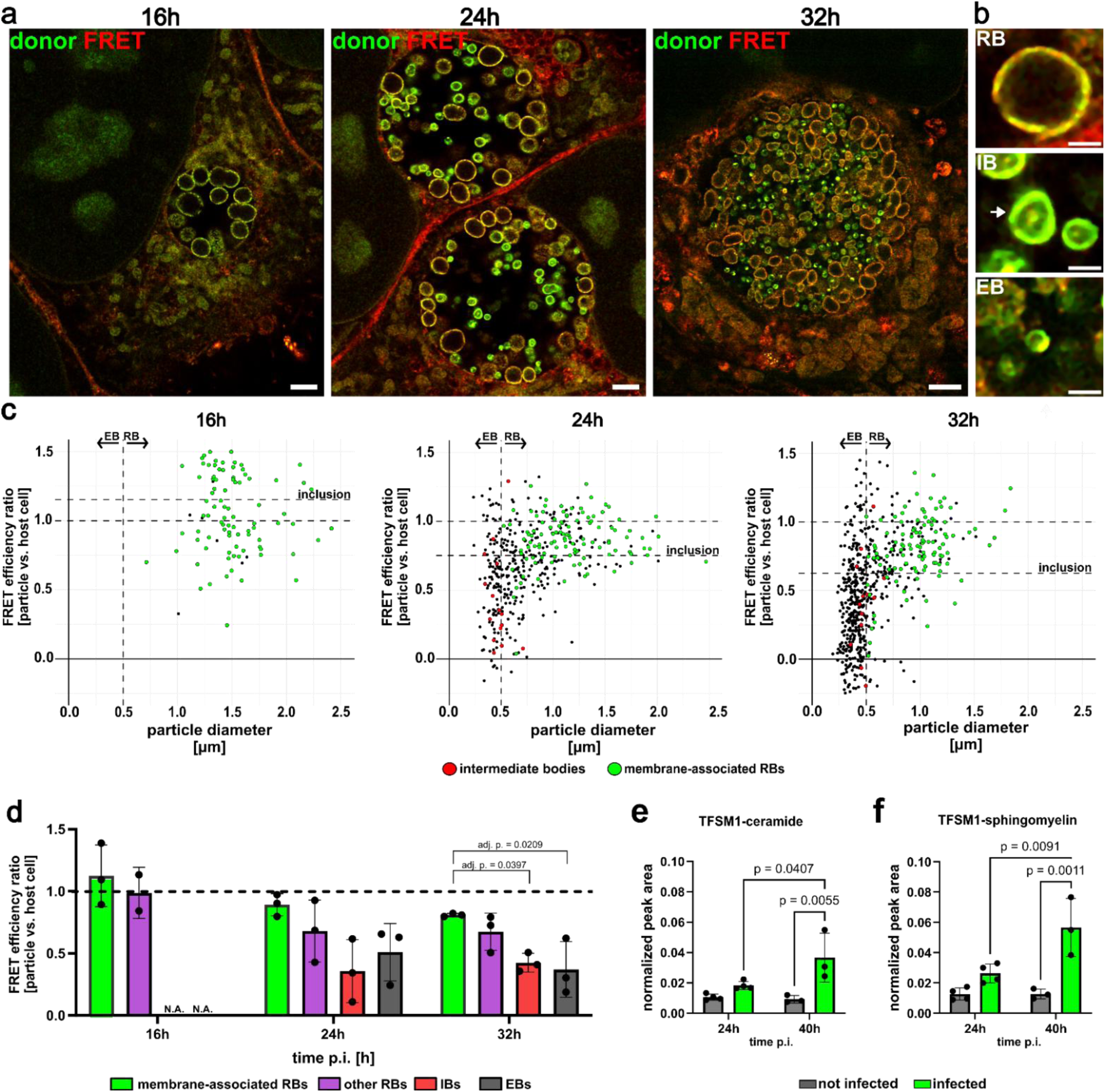
TFSM is metabolized within chlamydial inclusions. **a, b, Visualization of individual bacterial particles by 4x ExM.** HeLa 229 cells were infected with *C. trachomatis* in presence of TFSM2 and recorded via 4x ExM (a) at the indicated infection time. Chlamydial particles were classified as reticulate body (RB), intermediate body (IB) or elementary body (EB). Scale bar: 8 µm (∼2 µm with 4x expansion factor) in a, 2 µm (∼0.5 µm with 4x expansion factor) in b. **c, 4x ExM enables measuring the TFSM content of individual chlamydial particles.** Individual bacteria in inclusions recorded in a, were selected. These particles were classified as RBs associated with the inclusion membrane, “other” RBs (RBs not-associated with the inclusion membrane), intermediate bodies (IB, possess a characteristic TFSM accumulation in their center) and elementary body (EB). FRET efficiency in whole inclusion (horizontal line) and individual particles (dots) were determined and normalized to the FRET efficiency measured in a host cell area (set to 1). Normalized FRET efficiency (ratio “particle vs. host cell”) was plotted against particle diameter. Particles smaller than 0.5 µm were considered as EBs. IBs are highlighted in red, RBs (particles >0.5 µm) that associate with the inclusion membrane are marked in green. Other particles are marked black. The average FRET efficiency detected in whole inclusions is indicated as dotted line. n=3. **d, Membrane-associated RBs possess higher FRET efficiency than IBs and EBs.** Individual particle classes were selected, and the average FRET efficiency was determined. n=3. **e, f, Untargeted HPLC-MS/MS screening for TFSM metabolites.** HeLa 229 cells were infected with *C. trachomatis* or left uninfected in presence of TFSM1. After 24 h or 40 h p.i. lipids were extracted and analyzed by HPLC-MS/MS. Peak areas of TFSM1-ceramide (e) and TFSM1-SM (f) and were normalized to the sum of C16:0 SM and C16:0 ceramide peaks as a measure of the amount of cellular material used for lipid extraction. n ≥3. Statistics: One-way ANOVA and Tukey’s multiple comparison (d), two-way ANOVA and Tukey’s multiple comparison (e, f).

To further resolve the dynamics of TFSM metabolization on single particle level, we determined the FRET efficiency of whole inclusions as well as of individual bacteria by acceptor bleaching and normalized the results to the FRET efficiency detected in host cell areas (**Fig. 6**, c). We plotted the diameter of the particles against the corresponding FRET efficiency (**Fig. 6**, c) to distinguish between RBs (diameter > 500 nm) and EBs (diameter < 500 nm) and further identified RBs associated with the inclusion membrane (“membrane RBs”). A small subset of particles exhibited a characteristic TFSM accumulation in their center and we categorized these bacteria as intermediate bodies (IBs, **Fig. 6**, b).

At 16 h p.i., we exclusively detected RBs that were predominantly associated with the inclusion membrane and possessed similar TFSM content as the host cell (**Fig. 6**, c and d). TFSM content detected in membrane-associated RBs slightly decreased during infection (24 h p.i. and 32 h p.i.) but remained similar compared to host cell areas. This indicates a similar TFSM content of membrane-associated RBs and host cells also at later stages of infection. By contrast, we measured a significantly decreased FRET efficiency in IBs and EBs, suggesting a higher proportion of metabolized TFSM contained by these particles. In summary, our data suggest that membrane-associated RBs predominantly take up native TFSM, which is metabolized during RB-to-EB differentiation (**Fig. 6**, d) and subsequently accumulates on the nucleoid-containing EB cell pole (**Fig. 1**, **Fig. 2**).

### SM is formed from TFSM-derived ceramide during *C. trachomatis* infection

Although the FRET-based readout allows detection of TFSM metabolization with high spatial resolution, it does not provide the chemical identity of TFSM metabolite(s). To identify TFSM-derived metabolites formed during infection, we infected HeLa 229 cells in the presence of TFSM1, extracted the lipids, and screened for predefined TFSM1 metabolites by high pressure liquid chromatography coupled to tandem-mass spectrometry (HPLC-MS/MS; **Supp. Fig. 8**). Among the molecules that we included in the screen, only the TFSM1-ceramide and, surprisingly, the TFSM1-SM metabolites were found in detectable concentrations (**Fig. 6**, e and f).

Importantly, TFSM1-SM contains the functionalized amine and azide groups of TFSM1 in the backbone and the acyl chain but possesses a natural phosphocholine head group as confirmed by accurate mass and MS/MS fragmentation patterns. Interestingly, levels of both metabolites increased between 24 and 40 h p.i. and were markedly higher than in uninfected control cells during late-stage infection (**Fig. 6**, f). This increase was more pronounced for the TFSM1-SM metabolite compared to TFSM1-Cer. These observations support a model in which TFSM is transiently metabolized to ceramide, that subsequently serves as a substrate for TFSM1-SM synthesis, resulting in the accumulation of TFSM1-SM in the inclusion. This model would also explain the apparent discrepancy discussed earlier, as TFSM1-SM in contrast to TFSM1-Cer, possess a polar headgroup, which could interact with DNA in the nucleoid. ^64–71, 74^ Consequently, TFSM1-SM represents a plausible candidate for the sphingolipid species associated with condensed chlamydial nucleoids.

In summary, we propose a model in which SM is acquired by RBs attached to the inclusion membrane (**Fig. 7**). During RB-to-EB redifferentiation, SM is transiently metabolized to ceramide and then resynthesized to SM, which is then incorporated into to the bacterial nucleoid. EBs are released by host cells and subsequently infect neighboring cells. Upon host cell entry, the lipid-DNA association is resolved and DNA decondensation is initiated.

**Fig. 7.**
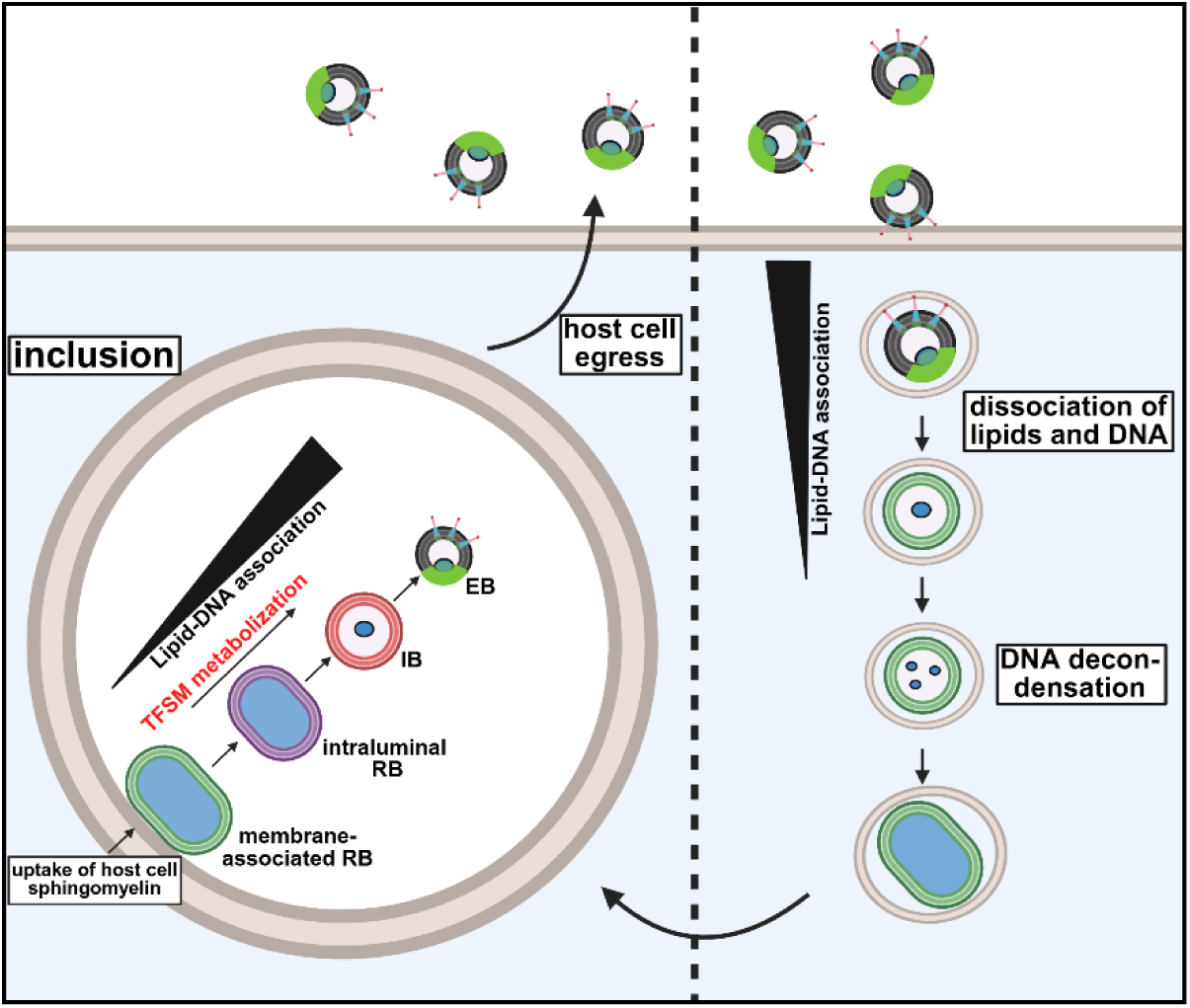
Proposed model for the role of sphingolipids during *C. trachomatis* infection. Sphingomyelin (SM) is taken up from the host by membrane-associated RBs. During RB-EB redifferentiation, SM is metabolized to ceramide (Cer) and resynthesized to SM, which is incorporated in EB nucleoids. EBs are released and internalized by neighboring cells. During RB-EB differentiation, first lipids are released from nucleoids and DNA decondensation is initiated thereafter.

## Discussion

Despite intensive research, the role of lipids in DNA organisation remains poorly understood, largely due to the limited methodology for studying the interactions between these two molecule classes.^75^ The presence of lipid converting enzymes^76–78^ and lipid droplets ^79–81^ in eukaryotic nuclei strongly suggests that lipids can modulate genetic information. However, only a few examples linking lipids directly to DNA regulation have been described. For instance, the bioactive lipid sphingosine-1-phosphate can directly impact epigenetics by interacting with histone deacetylases^82^ and SM-enriched membrane domains can influence transcription.^83^ Cholesterol has been shown to bind eukaryotic histones and thus, it is incorporated into nucleosomes.^84–86^ In bacteria, DNA is tethered to the cell envelope, which was hypothesized to facilitate the co-translational membrane incorporation of transmembrane proteins.^87–92^

Here, by combining ExM and cryo-ET, we delineate the lipid distribution in chlamydial EBs and uncovered a massive colocalization of lipid derivatives with the nucleoids of these particles as ∼90% of the nucleoid area contained the lipid analogues in our measurements (**Fig. 3**, h). In contrast, membranes labeled with pGk13a exhibited a more variable spatial relationship with the nucleoid. In some EBs, the pGk13a signal only partially overlapped with the nucleoid (**Supp. Fig. 4**, c and d), whereas in others, an extensive overlap between DNA and membrane signal was observed (**Supp. Fig. 4,** e and f). Moreover, pGk13a did not reveal the characteristic globe-shape lipid protrusion into the area containing the nucleoid as found for samples stained by TFSM. The interpretation of these observations is limited by the spatial resolution of the current approach. Clear visualization of the interface between membranes and DNA lays beyond the resolution limit of 8x ExM, which, on a conventional confocal microscope, is restricted to ∼30 nm. Hence, it remains elusive whether the nucleoid in these particles is in close proximity to the membrane or if membranes also penetrate into the membrane-facing side of the nucleoid.

One possible explanation for the differences between TFSM and pGk13a labeling is that these probes visualize distinct lipid pools. While pGk13a was designed to stain membranes, it likely does not label lipids bound to proteins or DNA. This could explain the discrepancy between the pGk13a and TFSM labeling, as latter could form complexes with protein or DNA thereby resulting in a distribution throughout the whole nucleoid.

Importantly, the lipid derivatives used for ExM possess C_6_ acyl chains, which are shorter than those found in naturally occurring lipids where fatty acid residues usually contain 14 or more C atoms.^93, 94^ Acyl chain length affects the properties of lipid molecules, for instance, lipid analogs containing shorter chain lengths are more likely to undergo spontaneous intermembrane transfer in liposome-based *in vitro* systems.^95–97^ Thus, TFSMs transport and localization could differ from natural long chain SM species. Our current methodology does not allow the visualization of natural lipids or long chain lipid analogs via ExM. In future studies, it will be important to develop approaches that enable visualization of naturally occurring lipids at nanoscale resolution to investigate whether these lipids are as well incorporated into EB nucleoids.

Although direct interaction of DNA with SM^74^ and other zwitterionic lipids ^64–71^ has been demonstrated *in vitro*, a structural role of phospholipids in DNA organization within a living organism has, to our knowledge, not been reported. The use of lipids for DNA condensation could have important benefits for an obligate intracellular bacterium. First, because lipids are hijacked from the host cell, this strategy may present a more energy-efficient alternative to *de novo* synthesis of histone-like proteins. Still, it remains elusive whether, and if so, how lipids can actively contribute to DNA condensation in *C. trachomatis*. Interestingly, a recent study reported that nucleoid formation is independent from histone-like proteins.^98^ Hence, it is tempting to speculate that lipids can functionally replace protein-mediated DNA condensation. In addition, dynamic remodeling of lipid–DNA interactions through lipid metabolism or lipid exchange could provide a mechanism for the rapid initiation of nucleoid decondensation and, consequently, the regulation of genome accessibility during EB-to-RB differentiation.

Consistent with this notion, dissociation of TFSM from nucleoids is one of the earliest events observed during DNA decondensation and occurs shortly after EBs enter their host cell and before the nucleoid decondenses (**Fig. 4**, b). Other studies have reported expression of genes in *C. trachomatis*^99^ and other *Chlamydia*^100, 101^ at this early time point during infection, which might be initiated through dissociation of the nucleoid from membranes.

The stimulus for and the mechanism behind DNA unwrapping in EBs is mostly unknown and to date only a methylerythritol phosphate metabolite has been described to disrupt DNA-histone interactions.^26^ Interestingly, we measured uptake of TFSM2 by EBs attached to the plasma membrane and its subsequent incorporation into nucleoids (**Fig. 4**, d). This raises the possibility that EBs could also take up negatively charged lipids such as phosphatidyl serine or phosphoinositides, the latter of which are enriched in phagosomal membranes.^102^ Alternatively, *C. trachomatis* encodes several D-type phospholipases with unknown function^103^, which could convert zwitterionic to negatively charged phospholipids. In both scenarios, presence of negatively charged lipids in nucleoids could destabilize the DNA complex and initiate DNA decondensation.

Important previous studies investigated early events after host cell entry of *C. trachomatis*^99^ and other *Chlamydia*^104, 105^ by electron microscopy (EM). However, DNA and lipids cannot be labeled specifically in EM, and localizing nucleoids is further complicated by a general high electron density of EBs upon negative staining and the limited image quality of older micrographs. Similar challenges exist in cryo-ET, as we and others^27^ only indirectly identified nucleoids by detecting the ribosome-depleted areas within EBs. Consequently, both the incorporation of lipids into nucleoids as well as the well-orchestrated DNA decondensation processes during EB-to-RB differentiation have likely been overlooked in previous studies. In contrast, 8x ExM enables fluorescent labeling of DNA and lipid derivatives with ∼30 nm spatial resolution, allowing precise three-dimensional tracking of DNA decondensation in particularly small EBs (**Fig. 4**, b, c). Using ExM, we show that EB-to-RB differentiation proceeds through multiple morphologically distinct intermediate forms (**Fig. 4**, b, c), which can be distinguished by the structure and organization of the DNA within the particles. Importantly, dissociation of TFSM from and detachment of the nucleoid from the inner bacterial membrane are very early events in EB-to-RB differentiation and were observed in intracellular bacteria already 1 h p.i.. Thereafter, DNA is decondensed stepwise, where the nucleoids decompose into several DNA foci between 4 and 5 h p.i. and are completely decondensed 6 h p.i..

Furthermore, our study provides new insights into sphingolipid metabolism within chlamydial inclusions. This advance was made possible by establishing quantitative FRET measurements of individual bacteria, which is an important improvement of our previous methodology, where we determined TFSM content of bacteria by calculating FRET/donor ratio images.^45, 46^ By our advanced imaging approach, we identified that TFSM is metabolized during RB-to-EB redifferentiation (**Fig. 5**) and that TFSM metabolites accumulate in mature EBs (**Fig. 6**, c and d). This hypothesis is further supported by the impaired TFSM metabolization upon overexpression of the chlamydial sRNA CtrR3, which blocks RB-to-EB redifferentiation. However, overexpression of CtrR3 strongly alters the expression profile of *C. trachomatis*^73^, and therefore its effects are not limited to developmental regulation. Nevertheless, the pronounced effect of CtrR3 overexpression on TFSM metabolism may prove a useful tool for identification of sphingolipid-metabolizing enzymes expressed by *C. trachomatis* in future studies.

Surprisingly, we only measured enhanced levels of TFSM1-ceramide and TFSM1-SM upon infection by mass spectrometry, whereas other TFSM1 metabolites were not found in detectable amounts (**Supp. Fig. 8**). While these data demonstrate that TFSM is predominantly metabolized to TFSM1-ceramide and TFSM1-SM during infection, the formation of other low abundant TFSM metabolites, which fall below the detection limit of our lipidomics approach, cannot be excluded.

Interestingly, TFSM1-SM and its upstream metabolite TFSM-Cer increased during the course of infection (**Fig. 6**, e and f). Since lipidomics analysis was performed with whole cell lipid extracts, these measurements cannot provide information about the cellular localization of detected lipid metabolites. However, our FRET efficiency measurements indicate a strong accumulation of TFSM metabolites within inclusions but not host cell areas during infection (**Fig. 5**, a and b). Together, these data support a model in which SM is metabolized to ceramide after RBs detach from the inclusion membrane and redifferentiate into EBs, followed by re-synthesis of SM, which ultimately accumulates in mature EBs. Consistent with our findings, previous studies demonstrated that *C. trachomatis* can synthesize SM in absence of the host cell SM machinery. ^48, 49^ However, the upstream metabolic processes and the origin of the ceramide used as substrate for SM biosynthesis remained elusive, because either natural lipids or lipid derivatives tagged only in the backbone were detected in these studies. By contrast, TFSMs harbor molecular tags in the head group and the ceramide backbone, which enabled detection of TFSM-derived SM metabolites (TFSM-SM) in which the alkyne-containing artificial headgroup is replaced by a natural phosphocholine head group. This facilitated the identification of SM resynthesize from TFSM-derived ceramide during *C. trachomatis* infection. In a related study^106^, we recently demonstrated that this generation of TFSM2-SM from TFSM2 is independent from host SM synthases, suggesting the presence of a chlamydial enzyme catalyzing SM biosynthesis as suggested previously.^48, 49^

The functional reason for the transient ceramide generation remains unclear. We speculate that ceramide is required for a certain step during EB formation e.g. for membrane remodeling or for subcellular localization of the lipid. Unlike SM, which contains a polar head group, ceramide can flip between the membrane leaflets^107^ and thus, transient ceramide generation might mediate the translocation of the lipid through the membrane bilayer, which in case of SM would require the action of specific enzymes.

CERT is crucial for inclusion development, since infection, consistently with previous reports ^108, 109^, was impaired in CERT K.O. cells (**Supp.** **Fig. 6**, e). The interplay between CERT and the complex SM metabolization described in the present study remains elusive. Previously, it was suggested that CERT fuels SM biosynthesis on the inclusion membrane by the host SM synthesis machinery^43^ or within the inclusion itself by a putative chlamydial SM synthase.^48, 49^ However, it is unclear whether infection depends on CERT-mediated ceramide transport to the inclusions or solely on the presence of CERT on the inclusions membrane for instance to form ER-inclusions contact sites as suggested by others.^108^ We suggest a model in which the activity of a chlamydial SM synthase is sustained by both, ceramide taken up from the host and SM-derived ceramide generated within the inclusion.

Beyond their structural role in membranes, we hypothesis that lipids may also play non-canonical roles in DNA organization. Future studies should aim to resolve the structure of the DNA-lipid-histone interface and determine whether i.) lipids directly interact with DNA, ii.) are associated with DNA-binding proteins or iii.) rather mediate the tethering of the nucleoid to the bacterial cell envelope. The interaction between lipids and DNA is challenging to investigate and is hence understudied.^75^ We propose that combining fixable lipid derivatives with super resolution imaging represents a powerful tool to further elucidate the role of lipids in DNA biology.

## Acknowledgements

We thank Dagmar Heuer (Rober Koch Institute, Berlin) for providing CERT K.O. cells and Adriana Moldovan for valuable scientific discussion. We thank Janna Eilts for valuable advice regarding super resolution microscopy. Kathrin Stelzner, Arpita Mohanty and Adriana Moldovan provided primers. Daniel Herrmann’s technical support during the HPLC-MS/MS analyses is greatly appreciated. We thank Christian Häring and Guru Amudhan for providing access to the Typhoon RGB scanner and their advice. The Deutsche Forschungsgemeinschaft (DFG) funded the Leica TCS SP5 CLSM under project code 116162193 and the Leica Stellaris 5 CLSM under project code INST93/1159-1 FUGG. ScopeM at ETH Zürich is acknowledged for access to cryoEM instrumentation. This work was supported by the DFG within the research training group RTG2581 (M.S, B.K., J.S., T.R.), and the Collaborative Research Center SFB 1583 (T.R.). T.R. receives funding from the European Research Council (ERC) under the European Union’s Horizon 2020 research and innovation program Grant No. ERC-2018-ADG/NCI-CAD (T.R.). M.R. was supported by funds of the Bavarian State Ministry of Science and the Arts and the University of Würzburg to the Graduate School of Life Sciences (GSLS), University of Würzburg. This work was supported by the DFG FOR3004 SYNABS, SA829/19-1 to M.S. and S.S.. M.S. and S.S. are supported by European Research Council (ERC) funding under the European Union’s Horizon 2020 research and innovation program (grant agreement No. 835102). We would like to acknowledge the assistance of the Core Facility BioSupraMol supported by the DFG. Work in Zurich (G.L.W.) was supported by the Fondation Jean-Jacques et Felicia Lopez-Loreta and by the Swiss National Science Foundation (SNSF) Grant TMSGI3_226208. Figures were created in *BioRender*.

## Author contributions

M.R., F.W. and T.R. conceptualized the study. M.R., F.W. and T.R. wrote the manuscript. M.R., F.W., P.K. T.E., F.S., S.S., B.K., M.S., G. L. W. performed and analyzed biological experiments. L.K., J.F. and J.S. performed chemical synthesis.

All authors contributed to the article and approved the submitted version.

## Materials & Methods

### Functionalized lipids

Functionalized lipids used in this study are shown in **Table 1**.

**Table 1.** Lipid derivatives used in this study.

| Lipids | Source | Reference |
| --- | --- | --- |
| α-NH<sub>2</sub>-ω-N<sub>3</sub>-C<sub>6</sub>-ceramide | Jürgen Seibel, Julius-Maximilians-University Würzburg, Germany | 44 |
| Trifunctional-Sphingomyelin 1 |  | 45 |
| Trifunctional-Sphingomyelin 2 |  | 45 |

### Cell culture

HeLa 229 cells (ATCC^®^ CCL-2.1™), HeLa wildtype (WT; ATCC^®^ CCL-2™), CERT K.O.^109^ and ASM K.O. cells^110^ were cultured in *standard medium* [RPMI+GlutaMAX^TM^ medium (Gibco^TM^, Cat. No. 72400054) containing 10 % (v/v) heat-inactivated (56 °C for 30 min) fetal bovine serum (FBS, Sigma Aldrich. Cat. No. F7524)]. McCoy cells (ATCC^®^ CRL-1696^TM^) were cultured in McCoy’s 5A (Modified) medium (Gibco^TM^, Cat. No. 26600080) containing 10 % (v/v) heat-inactivated FBS. Standard tissue culture procedures were used to maintain the cells The cells were cultured in humidified atmosphere with 5 % CO_2_ (v/v) at 37 °C using standard tissue culture procedures.

### *Chlamydia* culture

The preparation of *Chlamydia* was adapted from a previous publication.^111^ The L2 /434/Bu (ATCC VR-902B) serovar and the L2/434/Bu (ATCC VR-902B) serovar transformed with CtrR3 of *Chlamydia trachomatis* were used for this study. HeLa 229 cells were infected with *Chlamydia trachomatis* at MOI 1, which was then expanded for 48 h. The L2 CtrR3 strain was cultured in presence of 500 ng/mL spectinomycin-dihydrochloride pentahydrate (Sigma, Cat No. S4014-5G). Glass beads (2.85–3.45 mm, Roth, Cat. No. A557.1) were used to disrupt and mechanically lyse the cells, followed by centrifugation of the cells for 10 min at 755 × *g* at 4 °C. To collect the bacteria, the supernatant was centrifuged for 30 min at 40,000 × *g* at 4 °C. 1 x sucrose–phosphate–glutamic acid (SPG) buffer [7.5 % sucrose (Roth, Cat. No. 4621.2), 0.052 % KH_2_PO_4_ (Roth, Cat. No. 3904.1), 0.122 % Na_2_HPO_4_ (Roth, Cat. No. P030.2), 0.072 % L-glutamine (Gibco, Cat. No. 25030081)] was used to wash the bacterial pellets and the centrifugation step was repeated. The chlamydial pellet was resuspended in 1 x SPG buffer and passed through G20 (B. Braun, Cat. No. 612-0141) and G18 (B. Braun, Cat. No. 612-0147) hollow needles to singularize the *Chlamydia*. The bacteria were stored at -80 °C. Titration was used to determine MOI 1 and PCR was used to confirm that the *Chlamydia* were free from *Mycoplasma* contamination.

### *Chlamydia* infection in presence of lipid derivatives to visualize inclusions in mid-late infection cycle

HeLa WT and ASM K.O. cells were cultured in RPMI + GlutaMAX^TM^ medium containing 2 % heat-inactivated FBS two days before they were needed for experiments. HeLa 229 and HeLa WT or CERT K.O. cells were seeded a day prior to the experiment in a 24-well plate (6 × 10^4^ cells per well) on cover slips. The next day, medium was exchanged with fresh *standard medium* and the cells were infected with *C. trachomatis* at MOI 1. HeLa WT and ASM K.O. cells were seeded the day of the experiment in a 24-well plate (2 × 10^5^ cells per well) on cover slips in RPMI + GlutaMAX^TM^ medium containing 1 % heat-inactivated FBS. Cells were allowed to adhere for 6 h. The medium was exchanged with fresh medium containing 1 % heat-inactivated FBS and the cells were infected with *C. trachomatis* at MOI 1. In case of TFSM2 treatment, 3 h after infection the medium was exchanged to RPMI + GlutaMAX^TM^ medium containing 1 % heat-inactivated FBS and 10 µM TFSM2. In case of α-NH_2_-ω-N_3_-C_6_-ceramide, 1 h before fixation the medium was exchanged to RPMI + GlutaMAX^TM^ medium containing 10 % heat-inactivated FBS and 10 µM α-NH_2_-ω-N_3_-C_6_-ceramide. For GW4869 treatment, 10 µM GW4869 (Biozol, Cat No. MCE-HY-19363) was added 3 h after infection together with the TFSM2. For the L2 CtrR3 strain CtrR3 overexpression was induced by addition of 100 ng/mL anhydrotetracycline-hydrochloride (AHT; Supelco, Cat. No. 37919-100MG-R) 3 h after infection. In case of the CtrR3 strain, medium containing fresh AHT and TFSM2 was exchanged every 16 h. At the indicated time points, the cells were washed 2 × with Dulbecco’s phosphate-buffered saline (DPBS; Gibco, Cat. No. 14190169) processed as described below.

### *Chlamydia* infection of TFSM2-preloaded HeLa 229 cells to visualize the first 6 h of infection

HeLa 229 cells were seeded a day prior to the experiment in a 24-well plate (6 × 10^4^ cells per well) on cover slips. After 3 h, the medium of the cells was exchanged to fresh medium containing 1 % heat-inactivated FBS and 10 µM TFSM2. The next day, the cells were infected with *C. trachomatis* at MOI 25 (5 min and 1 h time point) or at MOI 15 (2 h to 6 h time point) and centrifuged for 10 min at 188x*g* at RT. At the indicated time points, the cells were washed 2 × with DPBS fixed and processed as described below.

### Uptake of BODIPY™ FL C12-Sphingomyelin

HeLa WT or CERT K.O. cells were seeded a day prior to the experiment in a 24-well plate (6 × 10^4^ cells per well) on cover slips. The next day, medium was exchanged with fresh *standard medium* and the cells were infected with *C. trachomatis* at MOI 1. In case of BODIPY™ FL C12-Sphingomyelin treatment, 3 h after infection the medium was exchanged to RPMI + GlutaMAX^TM^ medium containing 1 % heat-inactivated FBS and 10 µM BODIPY™ FL C12-Sphingomyelin (Thermo Fisher, Cat. No. D7711). 40 h after infection, the cells were washed 2 × with DPBS (Gibco^TM^). Fixation and DNA staining were performed as described below.

### Fixation, click staining of lipids and antibody staining

After infections were performed as described above, samples were fixed with 4 % PFA in PBS (Morphisto, Cat. No. 11762.0100) containing 0.25 % glutaraldehyde (Sigma, Cat. No. 10333) for 30 min/RT. Samples were washed thrice with PBS, permeabilized with 0.2 % Triton X-100 (Thermo Fisher, Cat. No. 28314) in PBS for 20 min/RT and again washed thrice with PBS. Samples that were to be stained with antibodies were blocked with 10 % FBS in PBS for 1 h/RT. Azide functions in lipid analogs were stained with BODIPY-FL-DBCO (conventional/unexpanded: 2 µM, 4x ExM: 4 µM, 8x ExM: 20 µM; Jena Bioscience, Cat.No. CLK-040-05) in Hank’s buffered salt solution (HBSS) for 1 h/37 °C at 170 rpm. Samples were washed 5x with PBS. Subsequently, alkyne groups of TFSM derivatives were stained with AZDye546 azide [Jena Bioscience, Cat. No. CLK-1283; product discontinued and we recommend using AZDye546-picolyl-azide (Jena Bioscience, Cat. No. CLK-1284) alternatively] for 1 h/37 °C at 170 rpm. The dye concentrations was depending on whether samples were expanded or imaged conventionally (conventional/unexpanded: 10 µM, 4x ExM: 20 µM, 8x ExM: 40 µM). Samples were washed 5x with PBS for 15 min on a see-saw rocker.

For antibody staining, samples were first incubated with primary antibody [anti-CERT, 1:100 (ExM), 1:200 (unexpanded), Abcam Cat.No. ab72536] and anti-chlamydial HSP60 (1:400, Enzo Life Sciences, Cat. No. ALX-804-071)] in 10 % FBS in PBS for 1 h/RT or at 4 °C overnight. Samples were washed thrice with PBS and incubated in 1:200 secondary antibody [goat anti-mouse Alexa Fluor 488 (ThermoFisher, Cat. No. A11001) or goat anti-rabbit CF568 (Sigma, Cat. No. SAB4600310)] in 10 % FBS in PBS for 1 h/RT or at 4 °C overnight. Samples were washed thrice in PBS. If indicated, unexpanded samples were stained with 5 µg/mL Hoechst 34580 (Thermo Fisher, Cat. No.H21486) in PBS for 30 min/RT and washed thrice with PBS for 15 min/RT on a see-saw rocker. DNA in ExM samples was stained after the expansion process (see below).

Subsequently, cover slips were either mounted in Mowiol [24g glycerol (Roth, Cat. No. 3783.2), 9.6 g Mowiol^®^4-88 (Roth, Cat. No 0713.2), 48 mL 0.2 M TRIS-HCl pH 8.5 (Sigma Aldrich, Cat. No T1503), 24 mL Millipore water] or further processed for ExM.

### Expansion microscopy

The protocols for 4x and 8x ExM are based on previous publications.^54, 55, 112, 113^ After antibody and/or click labeling, samples were postfixed with 0.25 % glutaraldehyde in PBS for 10 min/RT and washed thrice with PBS. For each specimen, a tube containing 96 µL 4x monomer solution [1x PBS (Sigma, Cat. No. D1408), 2.5 % (v/v) acrylamide (Sigma, Cat. No. A9926 or Sigma, Cat. No. A4058), 0,15 % (v/v) *N,N*’-methylenebisacrylamide (Sigma, Cat. No. M1533), 8.625 % (w/v) sodium acrylate (Sigma, Cat. No. 408220 or Santa Cruz, Cat. No. 236893), 2M NaCl (Sigma, Cat. No. S5886) in MilliQ water] or containing 97 µL 8x monomer solution [1xPBS, 1,1 M sodium acrylate, 2 M acrylamide (Sigma, Cat. No. A4058), 0.009 % (v/v) *N,N*’-methylenebisacrylamide in MilliQ water] were prepared. The cover slip was placed upside down into a drop (∼20 µL) of the respective monomer solution to remove residual PBS. Subsequently, 2 µL 10 % (w/v) ammonium persulfate (APS; Sigma, Cat. No. A3678) and 2 µL *N,N,N*’*,N*’-tetramethylethylenediamine (TEMED, Sigma, T9281) were added to tubes containing 4x monomer solution, while 1.5 µL 10 % (w/v) APS or 1.5 µL 10 % (v/v) TEMED were added to 8x monomer solution. Immediately, the solution was mixed and 56 µL (Ø 12 mm cover slip) or 80 µL (Ø 15 mm cover slip) were transferred on parafilm and the cover slip was placed into the drop upside down. Samples for 4x ExM were incubated for 75 min at RT in an enclosed chamber. Samples for 8x ExM were incubated for 1 h at 37 °C in an enclosed and humidified chamber. Afterwards, gels were removed from the cover slip and incubated with 8 U/mL proteinase K (Sigma, Cat. No. P4850) in digestion buffer [50 mM Tris-HCl pH 8 (Sigma, Cat. No. T1503), 1 mM EDTA (Sigma, Cat. No. ED2P), 0.5 % (v/v) Triton-X 100 (Thermo Fisher, Cat. No. 28314), 0.8 M guanidine HCl (Sigma, Cat. No. 50933) in MilliQ water] for 30 min/RT. Samples were incubated in excess of MilliQ water in Ø150 mm petri dishes and water was exchanged (usually 3-4 times) until samples were completely expanded.

Gels were cut into pieces of ∼1 cm x 1 cm and either stain for proteins (pan-ExM) and/or DNA or directly imaged in a humidified poly-L-lysin-coated (Sigma, Cat. No. A-005-C) imaging chamber (Ibidi, Cat. No. 82107) or on a silanized Ø24 mm cover slip in a sample holder (Thermo Fisher, Cat. No. A7816) flooded with MilliQ water.

### Pan-ExM and DNA staining of expanded samples

Expanded gels were cut into pieces of ∼2 cm x 2 cm and incubated in 40 µg/mL Atto594-NHS ester (pan-ExM^57^; Atto-Tec, Cat. No. AD594-31 or Sigma, Cat. No. 08741) and/or 12.5 µg/mL Hoechst 34580 (DNA staining; Thermo Fisher, Cat. No.H21486) in 100 mM NaHCO_3_ for 1.5 h/RT on a see-saw rocker. Afterwards, gels were washed 6-times with MilliQ water for 15 min each on a see-saw rocker at RT. Samples were imaged in a humidified poly-L-lysin-coated imaging chamber or on a silanized Ø24 mm cover slip in a sample holder (Thermo Fisher, Cat. No. A7816) flooded with MilliQ water.

### pGk13a staining and 8x ExM

The protocol for pGk13a staining was adapted from a previous publication.^61^ HeLa 229 cells were seeded a day prior to the experiment in a 24-well plate (6 × 10^4^ cells per well) on cover slips. The next day, medium was exchanged with fresh *standard medium* and the cells were infected with *C. trachomatis* at MOI 1. 32 h after infection, the cells were washed 2 x with DPBS. Cells were fixed with 4 % PFA supplemented with 0.5 % CaCl_2_ (Roth, Cat. No. 5239.1) in PBS overnight (overnight means ≥ 16 h) at 4 °C on a see-saw rocker. An 11.1 mg/ml pGk13a stock solution was prepared by dissolving pGk13a in 1:1 anhydrous DMSO (Sigma, Cat. No. 276855-1L):water (Thermo Fisher, Cat. No. J71786.AP). The samples were washed 3 x for 1 h at 4 °C with DPBS and afterwards were incubated in pGK13a labeling solution (98.5 % DPBS, 1.5 % pGK13a stock solution) overnight at 4 °C on a see-saw rocker. After quickly rinsing the samples with 1 x DPBS the samples were anchored using 0.1 mg/mL Acryloyl-X SE [6-((Acryloyl)amino)hexanoic acid, succinimidyl ester, Thermo Fisher, Cat. No. A20770] in DPBS overnight at 4 °C on a see-saw rocker. After the Acryloyl-X SE was discarded, samples were washed 3 x with DPBS for 1 h at 4 °C on a see-saw rocker. Next, gelation for 8x ExM and proteinase K digestion were performed as described above. The gels were transferred into 15 mL falcons each and were washed 3 x with DPBS for 30 min at RT on a roller mixer. Afterwards, DPBS was discarded, and the gels were incubated in 20 µM BODIPY-FL-DBCO in HBSS overnight at 4 °C on a roller mixer. The next day, the BODIPY-FL-DBCO solution was discarded, and the samples were washed 3 x with DPBS for 30 min at RT on a roller mixer. Samples were incubated in excess of MilliQ water in Ø150 mm petri dishes and water was exchanged (usually 3-4 times) until samples were completely expanded. Expanded gels were cut into pieces of ∼2 cm x 2 cm and DNA staining was performed as described above. Samples were imaged in a humidified poly-L-lysine-coated imaging chamber or on a silanized Ø24 mm cover slip in a sample holder flooded with MilliQ water.

### Cover slip silanization

Ø24 mm cover slips (VWR, Cat. No. 631-1583) were rinsed with MilliQ water and incubated with MilliQ water in an ultrasonic cleaning bath for 15 min. Subsequently, MilliQ water was exchanged with 5 M KOH (Roth, Cat. No. 6751.1) and cover slips were further incubated in an ultrasonic bath for 15 min. Cover slips were rinsed thrice with MilliQ water, incubated with 99 % EtOH (Sigma, Cat. No. 32221-2.5L-M) in an ultrasonic cleaning bath for 15 min and air dried. Each cover slip was covered with silanization solution [9 0% (v/v) EtOH, 5 % (v/v) AcOH (Roth, Cat. No. 3738.5), 5 % MilliQ water, 0.1 % (3-Aminopropyl)triethoxysilan (Sigma, A3648)], air dried and dipped in MilliQ water and 99 % EtOH until unbound silane was removed. Cover slips were stored at -20 °C.

### Determining TFSM metabolization by FRET acceptor bleaching

For unexpanded/conventional samples the in-built FRET AB wizard of the Leica TCS SP5 or Leica Stellaris 5 (Leica, Wetzlar, Germany) software was used. Areas containing an inclusion or a host cell membrane were selected for acceptor (AZDye546) bleaching and donor (BODIPY-FL) signals were determined pre- and post-bleaching. FRET efficiencies were calculated by the formular:

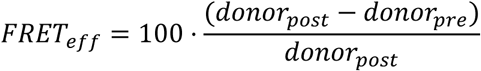

For each biological replicate, at least 5 different sites per sample were measured and results were averaged. Acceptor bleaching experiments of 4x ExM samples were performed at a Leica Stellaris 5 confocal microscope. Therefore, donor signal (BODIPY-FL) of infected cell(s) was recorded. Afterwards, the acceptor (AZDye546) was bleached within the whole frame, and the donor signal post-bleaching was recorded. Individual bacterial particles and host cell areas were selected manually in Fiji^114^ and FRET efficiency in all selected regions of interest (ROIs) were determined based on donor fluorescence pre- and post-bleaching. FRET efficiencies determined in individual particles were normalized to FRET efficiencies determined in host cell areas (set to 1) in the same image. Particle diameters were determined in Fiji and plotted against the corresponding FRET efficiency using ggplot^115^ and R studio.^116^ Particles were classified as “EBs” (particles with diameters <500 nm and not classified as IBs), “IBs” (particles with characteristic TFSM accumulation in their center, independent from particle diameter), “membrane-associated RBs” (particles with diameter >500 nm, attached to the inclusion membrane and not classified as IBs) and “other RBs” (particles with diameter >500 nm, not attached to the inclusion membrane and not classified as IBs). Negative FRET efficiencies were set to 0 and average normalized FRET efficiencies were calculated for each particle class.

### Analyzing EB nucleoids for lipid content and DNA condensation

HeLa 229 cells were infected with *C. trachomatis* for 32 h and processed for 8x ExM. Staining for TFSM2/α-NH_2_-ω-N_3_-C_6_-ceramide with BODIPY-FL-DBCO, for DNA with Hoechst 34580 and, if indicated, for pan-ExM with an Atto594 NHS ester was performed as described above. Then, regions within inclusions were imaged with a 63x oil objective and 3x optical zoom on a Leica Stellaris 5 confocal microscope. Images were further processed and analyzed in Fiji.^114^ For measuring the lipid content (BODIPY-FL signal) in individual nucleoids at mid/late-stage infection (here 32 h p.i.), regions containing RB DNA or host cell nuclei were removed from images. A median filter (2 pixels) was applied to the Hoechst channel, and nucleoids were selected with the “analyze particles” function in Fiji. Fluorescence mean intensities in all three channels of unprocessed images were measured and the average fluorescence of all measured nucleoids was determined using R studio. Samples not incubated with the clickable lipid derivatives but stained with BODIPY-FL-DBCO were used as controls. Ratios of samples incubated with lipid derivatives and untreated samples were calculated. For determining colocalization between lipid and DNA, BODIPY-FL channels were corrected for background staining. Therefore, the fluorescence mean intensity value detected in nucleoids of samples not incubated with the lipid derivatives were subtracted from all pixels of the BODIPY-FL channel. Then, a median filter (2 pixels) was applied to BODIPY-FL and Hoechst channels. RBs and EBs were analyzed separately. Therefore, all regions containing RBs were removed manually from the images for analyzing EBs and vice versa, all EBs were removed from images for RBs analysis. Subsequently, the colocalization between Hoechst and BODIPY-FL channels (Manders’ and Pearson’s coefficient) was determined with the Fiji plug-in “JACoP V2.1.4”.^117, 118^ Since EB nucleoids contained stronger BODIPY-FL signals compared to RB membranes, different thresholds were applied to RB and EB regions during determining Manders’ coefficient.

For analyzing DNA during early infection (5 min-6 h p.i.), extra- and intracellular regions were selected based on TFSM2 signal and a mask was created to categorize particles as intracellular or extracellular. Areas containing host cell nuclei were removed from the images. A median filter (2 pixels) was applied to the Hoechst channel and nucleoids were selected with the “analyze particles” function in Fiji. Size of the selected regions corresponds to the nucleoid area. Fluorescence mean intensities in all channels were measured in unprocessed images and the average fluorescence of all DNA particles was calculated in R studio. To correct background staining in the BODIPY-FL channel, the results of samples prepared in absence of TFSM2 were subtracted from TFSM2-treated samples, yielding the lipid content of nucleoids.

### Determining signal-to-background ratios

HeLa 229 cells were infected with *C. trachomatis* in presence or absence of TFSM for 32 h or 40 h. Samples were stained with BODIPYL-FL-DBCO and accumulations detected in EBs were selected manually using Fiji.^114^ Mean BODIPY-FL intensity in the selected areas were measured and the signal-to-background (STB) ratio was determined by the formular:

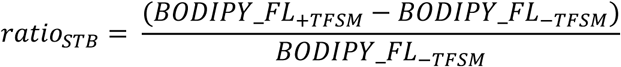

### Cryo-ET

#### Plunge freezing of C. trachomatis EBs

For cryo-ET, dense suspensions of *C. trachomatis* EBs (∼1’000’000 cells/µL), with or without TFSM treatment, were plunge frozen using a Vitrobot (Thermo Scientific) or a Leica GP2 (Leica Microsystems). Briefly, 3.5 μL of cell suspension was applied to glow-discharged Quantifoil R2/2 copper grids and blotted for 8 s only from the back, by replacing the front blotting paper with a Teflon sheet when the Vitrobot was used.^119^ The grids were then vitrified in liquid ethane–propane and stored in liquid nitrogen until further use.

#### Cryo-FIB milling

Electron-transparent lamellae were prepared following an established cryo-FIB workflow.^120^ Vitrified grids were clipped into AutoGrid rings (Thermo Scientific), transferred under cryogenic conditions to an EM VCM loading station (Leica Microsystems), where they were mounted onto a 40° pre-tilted cryo holder (Leica Microsystems). Using an EM VCT500 cryo-transfer system (Leica Microsystems), grids were transferred to an EM ACE600 (Leica Microsystems) and coated with ∼8 nm tungsten before transfer to a Crossbeam 500 cryo dual-beam FIB-SEM (Zeiss). Lamellae positions were selected using SEM imaging (3 - 5 kV, 58 pA), and milling patterns were defined and executed using SmartFIB. Rectangular patterns were milled with stepwise reduction of ion beam current (700, 300, 100 pA) and lamella thickness, followed by a final polishing step at 50 pA, targeting a final lamella thickness of ∼180 nm. Milled grids were stored in liquid nitrogen until imaging.

#### Cryo-ET data acquisition

Tilt series were acquired on a 300 kV Titan Krios G4 (Thermo Scientific) equipped with a K3 direct electron detector and a BioContinuum energy filter (20 eV slit, Gatan). Data were collected using SerialEM^121^ and SPACETomo^122^ at a pixel size of 2.678 Å, −6 μm defocus, and a total dose of ∼140 e⁻/Å². Dose-symmetric tilt series were recorded over ±60° with 3° increments. For **Fig. 3** and **Supp. Movie 2**, FIB-milled lamellae of EBs without TFSM were imaged, whereas for **Fig. 2**, plunge-frozen grids of EBs with and without TFSM were imaged without FIB milling.

#### Tomogram reconstruction, segmentation, and analysis

Tilt series were reconstructed using IMOD.^123, 124^ Frame alignment was performed with alignframes, followed by patch-tracking alignment and weighted back-projection at 4× binning. Tomograms were denoised and missing-wedge corrected using DeepDeWedge.^125^ For **Fig. 3**, b and **Supp. Mov. 1**, semantic segmentation was performed using a U-net trained in Dragonfly (Comet Technologies) on manually annotated slices, as described previously.^126^ Segmentations were visualized and smoothed in ChimeraX^127^ using the ArtiaXplugin.^128^

Subtomogram averaging of the type III secretion system (T3SS) for segmentation purposes was performed in Dynamo.^129^ T3SS particles were manually picked as dipoles, followed by extraction, alignment, and averaging. The resulting density was refined in ChimeraX and mapped back into the segmented volumes using ArtiaX.

For **Fig. 2**, tomograms of non-milled EBs were manually segmented in IMOD. Cells were approximated as spheres, T3SS positions were annotated, and membrane stacks were labelled in every tenth slice with smooth interpolation. Angles between T3SSs and membrane stacks were measured manually. Visualizations were generated in Blender (https://www.blender.org).

### Determining the orientation of the lipid accumulation and plasma membrane attachment

For measuring the orientation of the lipid accumulation during plasma membrane attachment of EBs, HeLa 229 cells preloaded with TFSM2 were infected with *C. trachomatis* for 1 h. TFSM2 was stained with click dyes as described above. Then, cells were visualized by 8x ExM, and z-stacks were recorded on a Leica Stellaris 5 confocal microscope. In Fiji, the orientation of the lipid accumulation in respect to the plasma membrane within individual EBs was analyzed. Therefore, a tangent was delineated manually on the EB-plasma membrane attachment site. Then, the angle between this tangent and the lipid accumulation within the EB was measured. Angles can take values between 0° and 90°.

For measuring the orientation between the T3SS and membrane stacks, an EB preparation was visualized by cryo-ET. The center of T3SS structures was determined manually and a vertical line between the T3SS center and the particle center was delineated. The same procedure was repeated for the center of the membrane stacks. The angle between the two lines describes the orientation of the T3SS in respect to the membrane stacks. Angles can take values between 0° and 180°.

### Measuring TFSM1 metabolites by lipidomics

HeLa 229 cells were seeded a day prior to the experiment in a 6-well plate (24 h time point: 5 × 10^5^ cells per well for infected samples and 2.5 × 10^5^ cells per well for uninfected samples, 40 h time point: 2.5 × 10^5^ cells per well for infected samples and 1.25 × 10^5^ cells per well for uninfected samples). The next day, the medium of the cells was exchanged with fresh *standard medium*. The cells were infected with *C. trachomatis* at MOI 1 or left uninfected. 3 h after infection, the medium was exchanged to medium containing 1 % heat-inactivated FBS and 10 µM TFSM1 for one well per time point and condition. At the indicated time points, cells were washed 2 × with DPBS, resuspended in 1 mL methanol (LC-MS CHROMASOLV, Fluka analytical, Cat. No. 34966-1L). Lipids were extracted as described^130^ and analyzed for the indicated TFSM1 metabolites (**Supp. Fig. 8**). Two strategies were used for this purpose. In an initial untargeted approach, the lipid extracts were separated chromatographically on a 1290 Infinity II HPLC (Agilent Technologies, Waldbronn, Germany) equipped with a Poroshell 120 EC-C8 column (3.0 × 150 mm, 2.7 μm; Agilent Technologies) and the TFSM1-derived lipid metabolites were identified using a 6550 quadrupole-time-of-flight (Q-TOF) instrument (Agilent Technologies). To confirm the structure of the TFSM1-Cer and TFSM1-SM metabolites identified in this way, they were further analyzed on a 6495C triple-quadrupole (QQQ) instrument (Agilent Technologies) with regard to their MS/MS fragmentation behavior. For both methods, analogous chromatographic conditions and, as far as possible, identical settings for the electrospray ion source were used. A previously published method for quantifying canonical cellular sphingolipids served as the basis.^130^ Metabolite identification was performed using MassHunter Qualitative Analysis software (version 10.0) with the “Find by Formula” algorithm. All detected metabolites had a mass error (Δ*m/z*) of less than 1 ppm. Peak assignment and integration were performed using MassHunter Quantitative Analysis software (for QTOF, version 10.1).

### Production of *C. trachomatis* CtrR3

Primers used for cloning procedures are listed in **Table 2**.

**Table 2.**
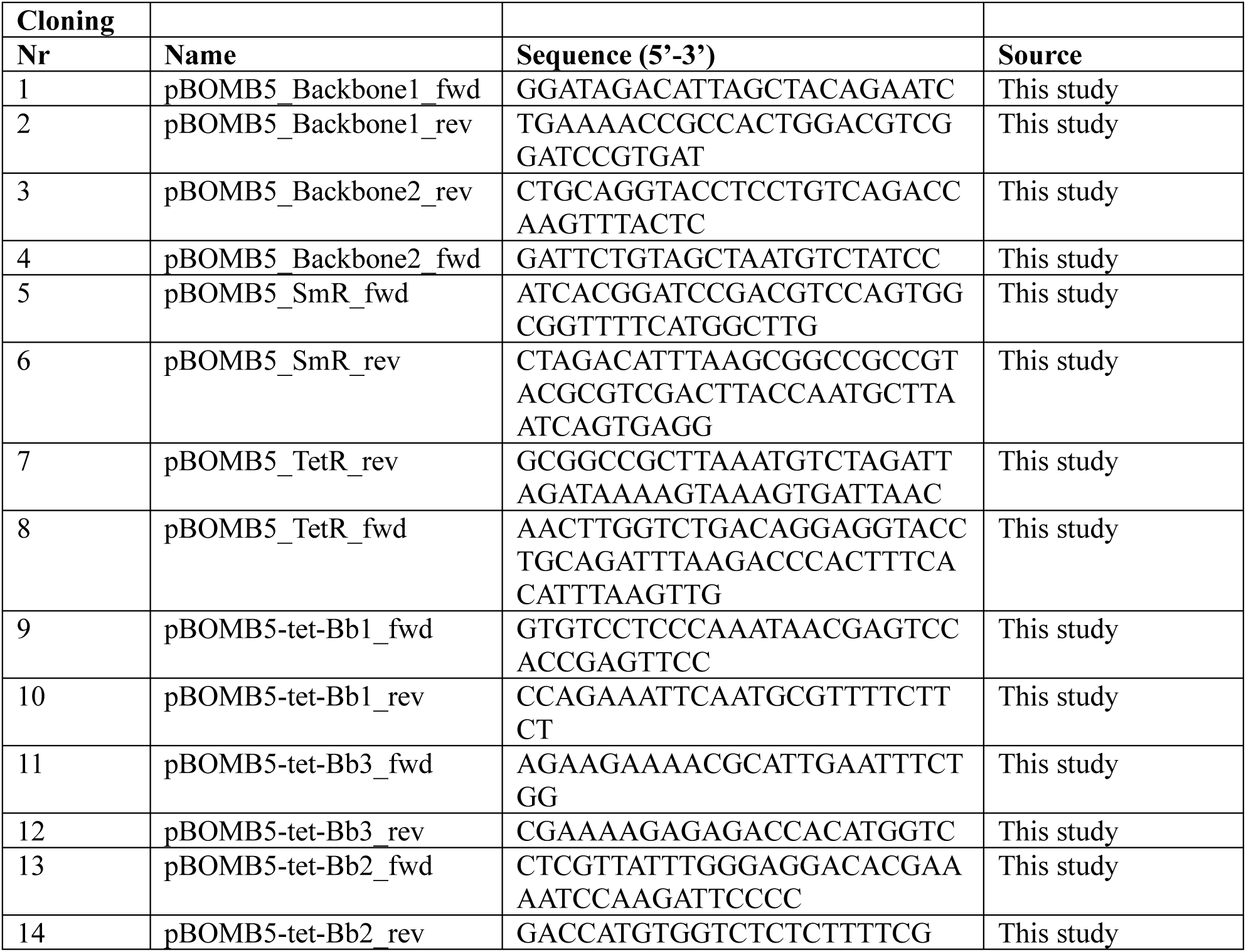

#### pBOMB5-tet-aadA-CtrR3 Plasmid Generation

The pBOMB5-tet-aadA plasmid was generated from the pBOMB4-tet-aadA plasmid in accordance with the already published model.^73^ DNA fragments were amplified using Phusion™ High-Fidelity DNA-Polymerase (ThermoFisher, Cat. No. F530) and PCR products were purified using the QIAquick PCR & Gel cleanup Kit (QIAGEN, Cat. No. 28506). *E. coli*-transformants were cultured in LB-Medium [10 % Peptone (Roth, Cat. No. 8952), 5 % yeast extract (Roth, Cat. No. 2363), 5 % NaCl (Roth, Cat. No. P029)] and LB-Agar plates [LB-Medium with 15 % Agar (Becton Dickinson, Cat. No. 257353)] containing 100 µg/mL spectinomycin. Plasmids were isolated using the NucleoSpin Plasmid Mini Kit (Macherey-Nagel, Cat. No. 740588) and NucleoBond Xtra Midi EF Midi Kit (Macherey-Nagel, Cat. No. 740420). The sequences of the final plasmids were confirmed by Sanger and Nanopore sequencing at Microsynth Seqlab GmbH. To generate the pBOMB5-tet-aadA plasmid, first the ampicillin resistance cassette and the tet-inducible expression cassette were removed, and the orientation of the Tet-Repressor (Tet-R) was reversed. To this end, the backbone was split into two fragments and amplified using the primers 1+2 and 3+4 respectively. The spectinomycin resistance cassette (SmR) was amplified using the primers 5+6, the Tet-R with the primers 7+8. The plasmid was generated using the In-Fusion Snap Assembly Master Mix (Takara, Cat. No. 638947) and transformed into Stellar competent cells (Takara, Cat. No. 636763).

Next, the CtrR3 sRNA backbone containing random bases in the hairpin was cloned into the pBOMB5-tet-aadA plasmid by digesting the pBOMB5-tet-aadA vector and a gBlock (Integrated DNA Technologies) encoding for the sRNA and the Tet-inducible promoter, with FastDigest SalI (Thermo Scientific, Cat. No. FD0644) and FastDigest NotI (Thermo Scientific, Cat. No. FD0594) and subsequently ligated using T4 DNA Ligase (Thermo Scientific, Cat. EL0011). The newly generated plasmid was transformed into chemically competent *E. coli* DH5α (homemade).

To change the bases in the hairpin of the CtrR3, the vector was amplified using the primer pairs 9+10, 11+12 and 13+14, followed by In-Fusion cloning and Stellar transformation. After successful sequence validation, the plasmid was transformed into chemically competent *E. coli* JM110^131^ (homemade) to obtain unmethylated plasmid DNA for *Chlamydia* transformation.

#### Chlamydia transformation

The pBOMB5-tet-aadA-CtrR3 plasmid was transformed into *C. trachomatis* serovar L2/434/Bu (ATCC VR-902B) as previously published.^132^ To prepare crude stocks of *C. trachomatis* for transformation, McCoy cells were infected with MOI 1. At 40 h p.i. the cells were lysed by scraping and the lysate was centrifuged at 20,000 x *g* and 4 °C for 30 min. The pellet was resuspended in appropriate amounts of 1xSPG. To pellet the cell debris, the suspension was centrifuged at 200 x *g* and 4 °C for 5 min. The crude stocks were stored at -80 °C. The MOI of the crude stocks was determined by performing a sham transformation. Therefore, McCoy cells were seeded in 6-well plates. The next day, different amounts of crude stock were mixed with 50 µL CaCl_2_-tranformation buffer [10mM Tris pH7.4 (Sigma Aldrich, Cat. No. T1503), 50 mM CaCl_2_ (Roth, Cat. No. 5239) in water], incubated for 30 min at RT and mixed every 10 min. The incubation was stopped by adding HBSS. This inoculum was used to infect the McCoy cells by replacing the medium in the wells followed by a centrifugation for 1 h at 900 x *g* and 20 °C. Afterwards, the HBSS was replaced with McCoy Medium + 10 % FBS + 1 µg/mL cycloheximide (Sigma Aldrich, 01810). MOI 1 was determined by light microscopy 24 h p.i. *C. trachomatis* was transformed using McCoy cells seeded in 6-well plates. For each well, 2 µg unmethylated plasmid DNA was mixed with 50 µL CaCl_2_-transformation buffer and *Chlamydia* of the crude stock to reach MOI 2. After 30 min incubation at RT with mixing every 10 minutes, the reaction was stopped by adding 2 ml HBSS and subsequently infecting McCoy cells (P0). After 1 h centrifugation at 900 x *g* and 20 °C the HBSS was replaced with McCoy + 10 % FBS + 1 µg/mL cycloheximide. Selection of the transformants was started 8 hpi by adding 500 µg/ml spectinomycin to the cell culture medium. After 48 hpi the infection was passaged onto fresh McCoy cells by lysing the infected cells with a cell scraper, centrifuging the lysate for 30 min at 20,000 x *g*/4 °C. The pellet was resuspended in 2 mL fresh McCoy Medium + 10 % FBS + 1 µg/mL cycloheximide + 500 µg/mL spectinomycin and centrifuged for 5 min at 200 x *g* and 4 °C. The supernatant was used to infect a fresh layer of McCoy cells by 15 min centrifugation of the plate at 900 x *g* and 37 °C This step was repeated after 48 h until fluorescent inclusions appeared at P2. Next, crude stocks were prepared, and chlamydia culture was performed as described above.

### ASM and NSM activity assays

The assays are based on previously published protocols.^133, 134^ HeLa 229 cells were seeded a day prior to the experiment in a 12-well plate (1 × 10^5^ cells per well for infection and 0.5×10^5^ for uninfected controls). The next day, the medium of the cells was exchanged with fresh *standard medium*. The cells were infected with *C. trachomatis* at MOI 1 or left uninfected for the indicated duration. Cells were transferred on ice, washed thrice with DPBS and 100 µL/well SMase lysis buffer [250 mM sucrose, 1 mM EDTA, 0.2 % TX-100, 1xRoche protein inhibitor cocktail (Sigma, Cat. No. 11873580001)] were added. Cells were scraped from the well, transferred into a reaction tube and incubated for 5 min/4 °C. Then, lysates were incubated in an ultrasonic bath for 5 min/4 °C and then further incubated for 5 min/4 °C. Lysates were cleared by centrifugation 20.000xg/4 °C/5 min. Protein concentrations were determined with a bicinchoninic acid (BCA) assay kit (Thermo Fisher, Cat. No. 23227) according to manufacturer’s instructions. For NSM assays, a volume corresponding to 10 µg protein was incubated with 100 µL NSM assay buffer [200 mM HEPES pH 7, 200 mM MgCl_2_, 0.05 % Nonidet P-40 (AppliChem, Cat. No. A1694,0250)] complemented with 0.58 µM BODIPY-FL-C_12_-SM (Thermo Fisher, Cat. No. D7711) for 22 h/37 °C/300 rpm. For ASM assays, a volume corresponding to 1 µg protein was incubated with 100 µL ASM assay buffer [500 µM ZnCl_2_, 200 mM sodium acetate pH 5.0, 500 mM NaCl, 0.02 % Nonidet P-40] complemented with 0.58 µM BODIPY-FL-C_12_-SM for 4h/37 °C/300 rpm.

Reactions were stopped by addition of 250 µL 2:1 CHCl_3_:MeOH. Samples were vortexed, centrifuged 13.000 x *g*/3 min/RT and 25 µL of the lower organic phase were transferred into a fresh reaction tube. The solvent was removed for 20 min/45 °C in a SpeedVac evaporator. The pellet was resuspended in 10 µL 2:1 CHCl_3_:MeOH and spotted in 2.5 µL steps onto a thin layer chromatography plate (Alugram, Xtra Sil G/UV254, layer 0.2mm/silica gel 60; Macherey-Nagel/VWR, Cat. No. 552-1006), which was developed with 80:20 CHCl_3_:MeOH. BODIPY-FL fluorescence was recorded with a Typhoon RGB scanner (Amersham).

Fluorescence intensities of the upper (BODIPY-FL-C_12_-ceramide, product) and lower band (BODIPY-FL-C_12_-SM, educt) were measured in Fiji. The enzyme activity was determined based on the amount of generated product, assay time and used protein amount.

### RT-qPCR

HeLa 229 cells were seeded a day prior to the experiment in a 6-well plate (2.5 × 10^5^ cells per well that will be infected and 1.25 × 10^5^ cells per well that will not be infected). The next day, the medium of the cells was exchanged with fresh *standard medium,* and the cells were infected with *C. trachomatis* at MOI 1 or left uninfected. For the L2 CtrR3 strain CtrR3 overexpression was induced 3 h after infection by addition of 100 ng/mL AHT. Medium containing fresh AHT was exchanged every 16 h. At the indicated time points, the cells were washed 2 × with DPBS and RNA was isolated using the RNeasy Mini Kit (Qiagen, Cat. No. 74104) according to manufacturer’s instructions. DNA was removed using the RNase-Free DNase Set (Qiagen, Cat. No. 79254) and cDNA was generated using the Revert Aid First Strand cDNA Synthesis Kit (Thermo Scientific, Cat. No K1622) according to manufacturer’s instructions. The GreenMasterMix (2X) High ROX (Gennaxon, Cat. No M3052.0500) and the StepOnePlus system (Life Technologies) were used to perform the RT-qPCR according to the StepOne software (version 2.3) and data was reanalyzed using the Design & Analysis Software 2.7.0 (Thermo Scientific). Analysis of the data was performed according to the 2^(−ΔΔCt)^ method^135^ using Excel (Microsoft). YWHAZ was used as endogenous control for human genes, OMCB was used as endogenous control for chlamydial genes. Primers used for RT-qPCR are shown in **Table 3**.

**Table 3.**
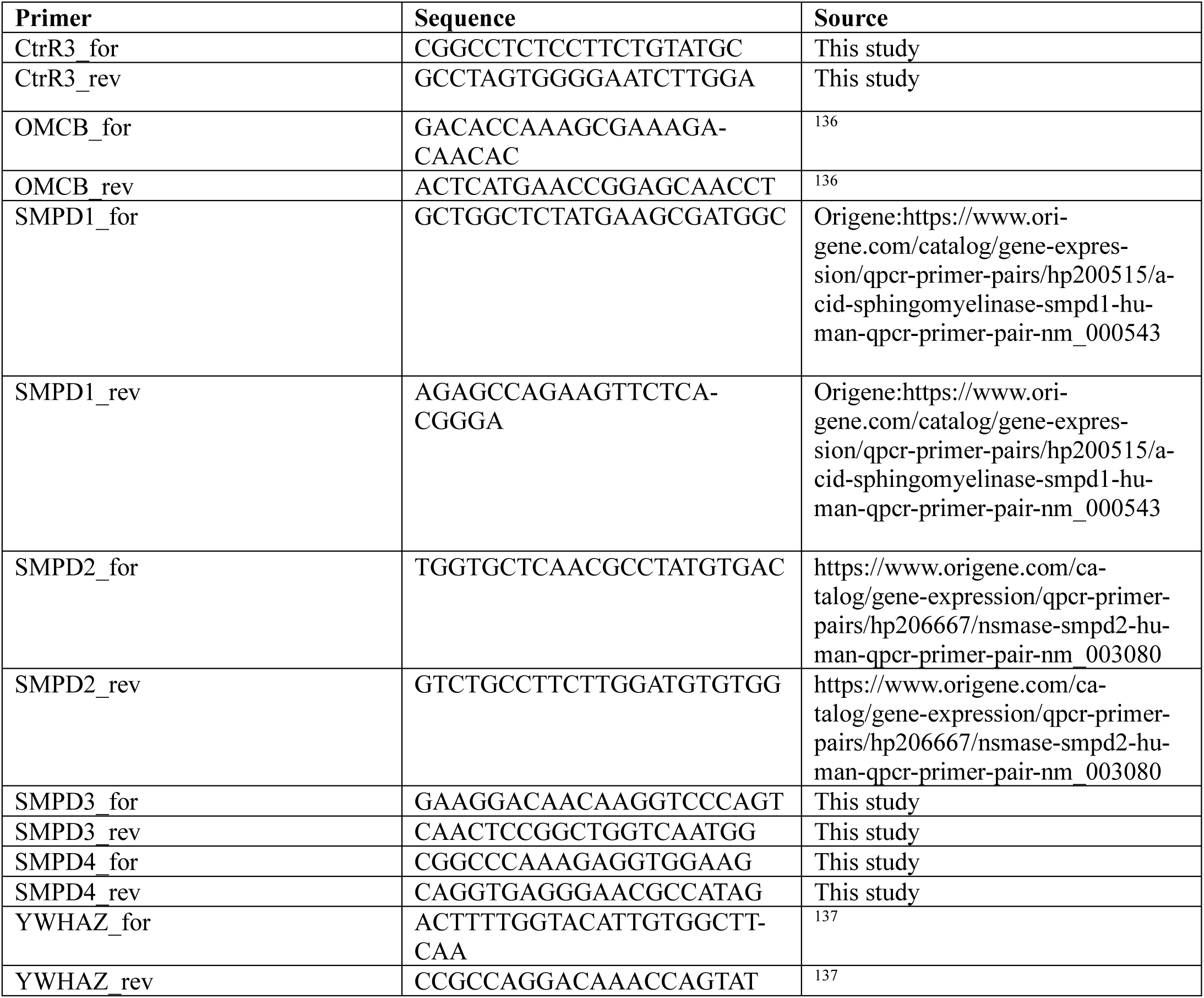

### Statistics and data representation

Statical analysis was performed using GraphPad Prism (Version 10.1.2. and Version 11.0.1). Details can be found in the corresponding figure legends. All data are represented as mean ± standard deviation.

## Supplementary Information

**Supp. Fig. 1.**
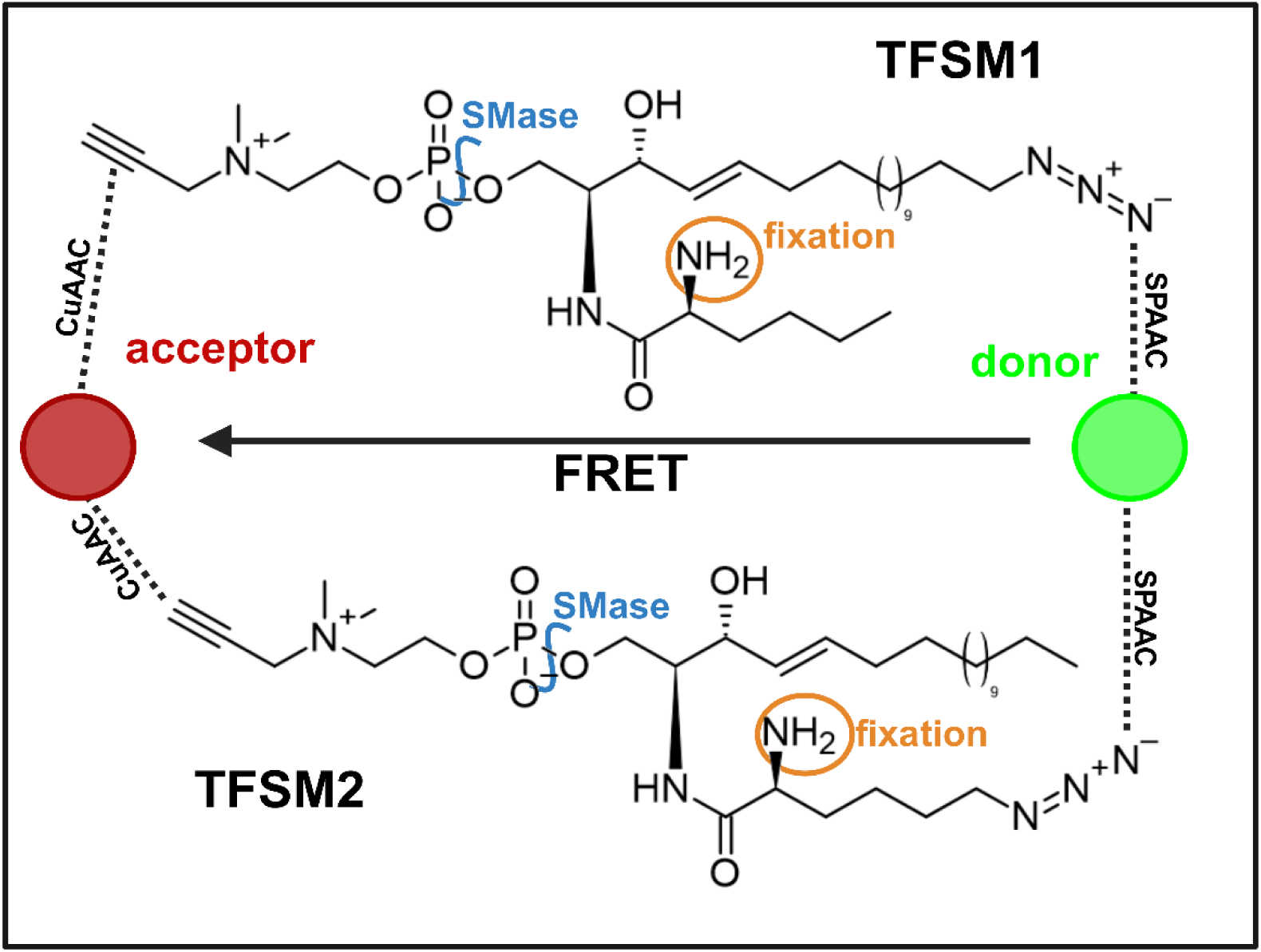
Principle of trifunctional sphingomyelin derivatives. Compared to naturally occurring SM, TFSMs possess an additional amino function (orange) for fixation and linkage into a swellable hydrogel for ExM. Both derivatives possess an alkyne in their head group, which can be modified with a fluorophore (red, acceptor) by copper catalysed azide-alkyne cycloaddition (CuAAC). TFSM1 harbours an azido function in the sphingoid backbone, TFSM2 in the fatty acid chain. Azido functions can be equipped with a fluorophore (green, donor) via strain-promoted azide-alkyne cycloaddition (SPAAC). Fluorophores clicked to the head group and the backbone can interacted via Förster resonance energy transfer (FRET). FRET can be used as readout for metabolization of the molecules, since the FRET system is destroyed upon cleavage by an SMase (blue).

**Supp. Fig. 2.**
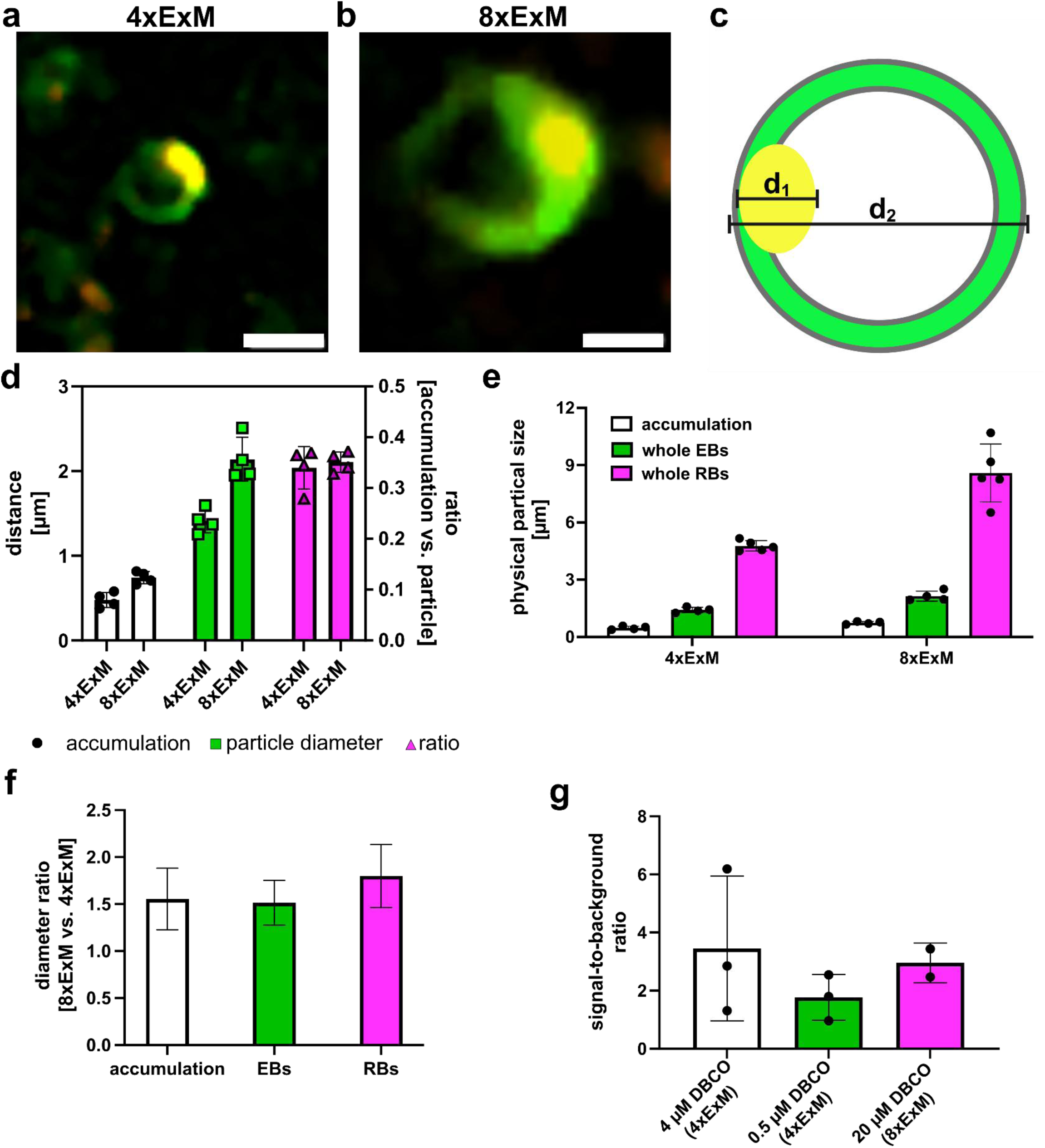
Lipid accumulation in EBs is not due to expansion or staining artifacts. HeLa 229 cells were infected with *C. trachomatis* in the presence or absence of TFSM. Samples were visualized by 4x or 8x ExM as indicated. **a-d, uniform expansion of EBs and their lipid accumulation.** Expansion of EBs upon 4x ExM (a) and 8x ExM (b) was compared. Therefore, the diameter of the lipid accumulation (d_1_) and whole EBs (d_2_) were measured, and ratios “accumulation vs. particle” were calculated (c, d). n =4. Scale bar: 1 µm (∼250 nm with 4-fold expansion factor [a] or ∼125 nm with 8-fold expansion factor [b]). **e, f, size comparison of bacterial particles upon 4x and 8x expansion.** Average diameters of RBs, EBs and the EB-associated lipid accumulation were measured in 4x and 8x expanded samples (e) and ratios “8x ExM vs. 4x ExM” of these structures were calculated. n≥4. (f). **g, determining signal-to-background ratios in lipid accumulations.** Fluorescence mean intensity (mean_FL_) in lipid accumulations was measured in samples incubated with and without TFSM (e). Signal-to-background ratios were determined by [mean_FL_(+TFSM) - mean_FL_(-TFSM)]/ mean_FL_(-TFSM).

**Supp. Fig. 3.**
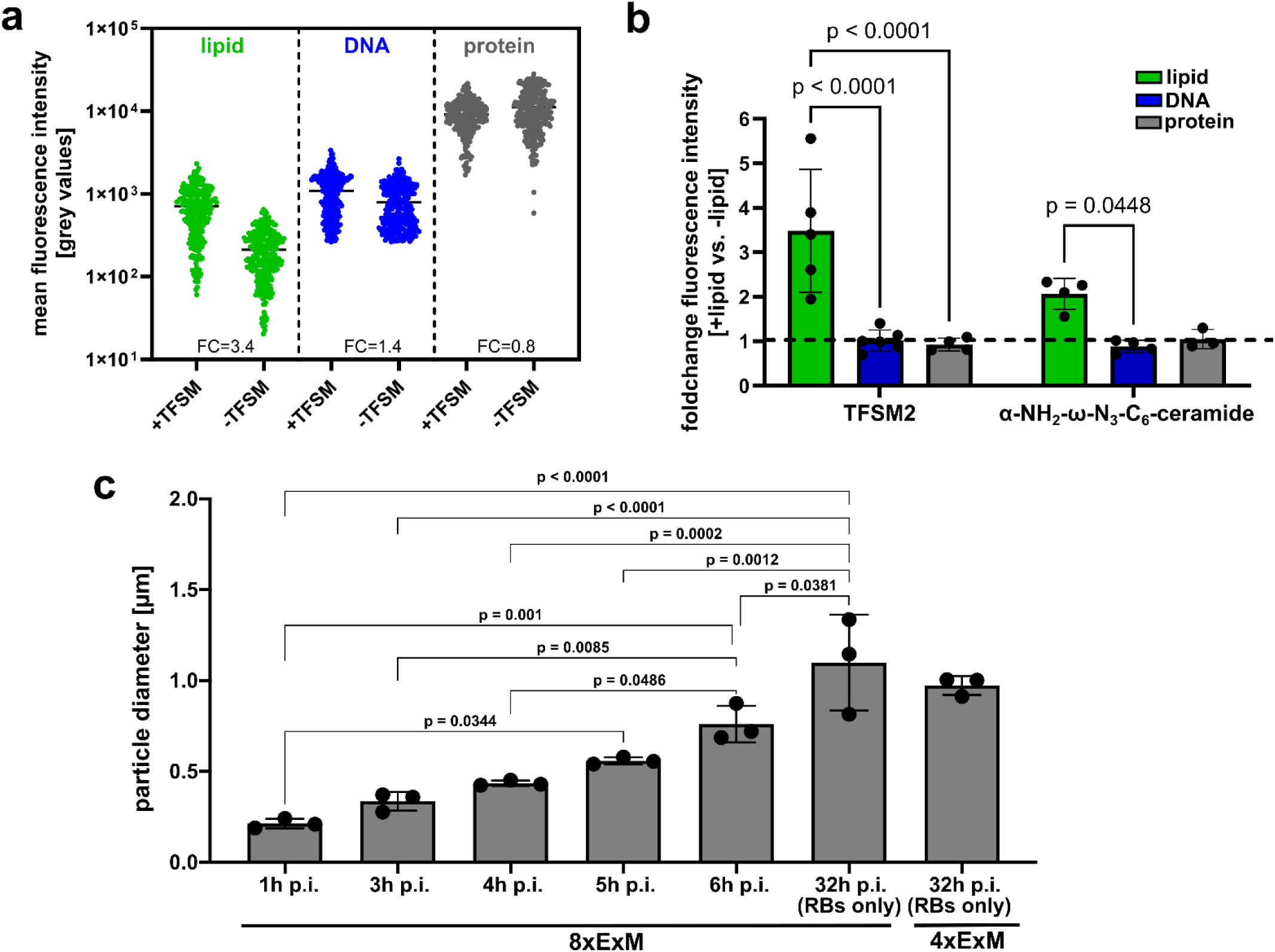
TFSM2 and α-NH_2_-ω-N_3_-C_6_-ceramide are enriched in chlamydial nucleoids. **a, b, quantification of TFSM2 and α-NH_2_-ω-N_3_-C_6_-ceramide in chlamydial nucleoids.** HeLa 229 cells were infected with *C. trachomati*s in the presence or absence of TFSM2 or α-NH_2_-ω-N_3_-C_6_-ceramide and imaged by 8x ExM. Regions containing EB nucleoids were selected in the Hoechst channel and fluorescence mean intensity in the lipid, the NHS ester and Hoechst channel were measured (a). The average fluorescence intensity in all detected nucleoids was determined in lipid, DNA and protein channels. Then the foldchange between samples infected in presence (+lipid) or in absence (-lipid) of the lipid analogues was determined in the respective channels. **c, particle size of intracellular bacteria continuously increases in the first 6 h of infection.** HeLa229 cells were infected in presence of TFSM, fixed at the indicated time points and visualized by 8x or 4x ExM. Particle diameters of intracellular bacteria were measured and corrected for the respective expansion factors (divided by 4 or 8). For 32 h p.i., only RBs (see Figure 5 for particle classification) were used for the measurement. Statistics: Mixed-effects analysis (b) or one-way ANOVA (c) and Tukey’s multiple comparison.

**Supp. Fig. 4.**
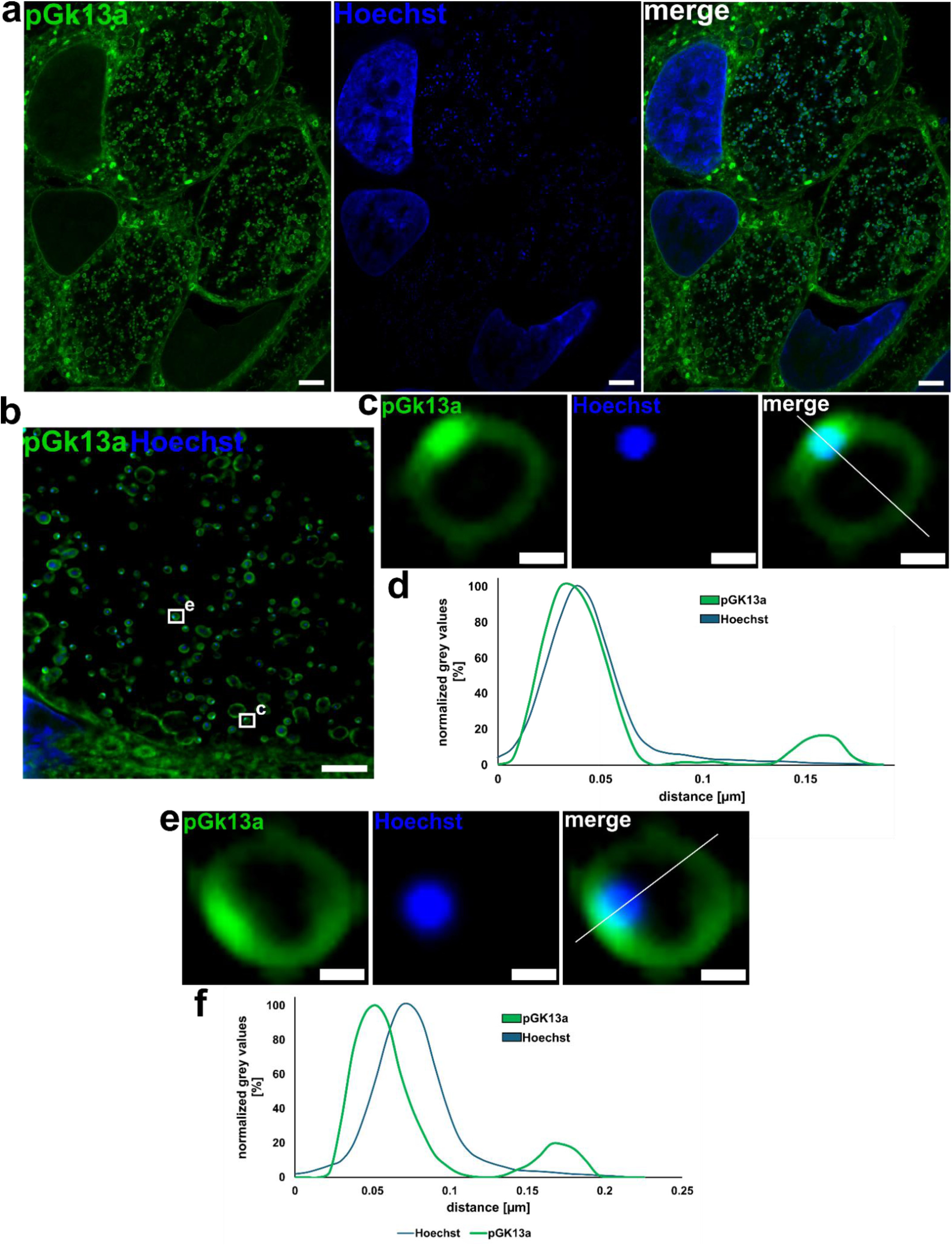
The EB nucleoid is in close association with the membrane accumulation. HeLa 229 cells were infected with *C. trachomatis* for 32 h, fixed, stained with pGk13a and 8-fold expanded. DNA was stained with Hoechst. Plot profiles of two individual EBs (c and g) were measured for pGk13a and Hoechst (d, f). Distances in plots were corrected for the expansion factor. Scale bars: a: 16 µm (∼2 µm with 8-fold expansion factor), b: 8 µm (∼0.5 µm with 8-fold expansion factor), c, e: 0.4 µm (∼ 50 nm with 8-fold expansion factor).

**Supp. Fig. 5.**
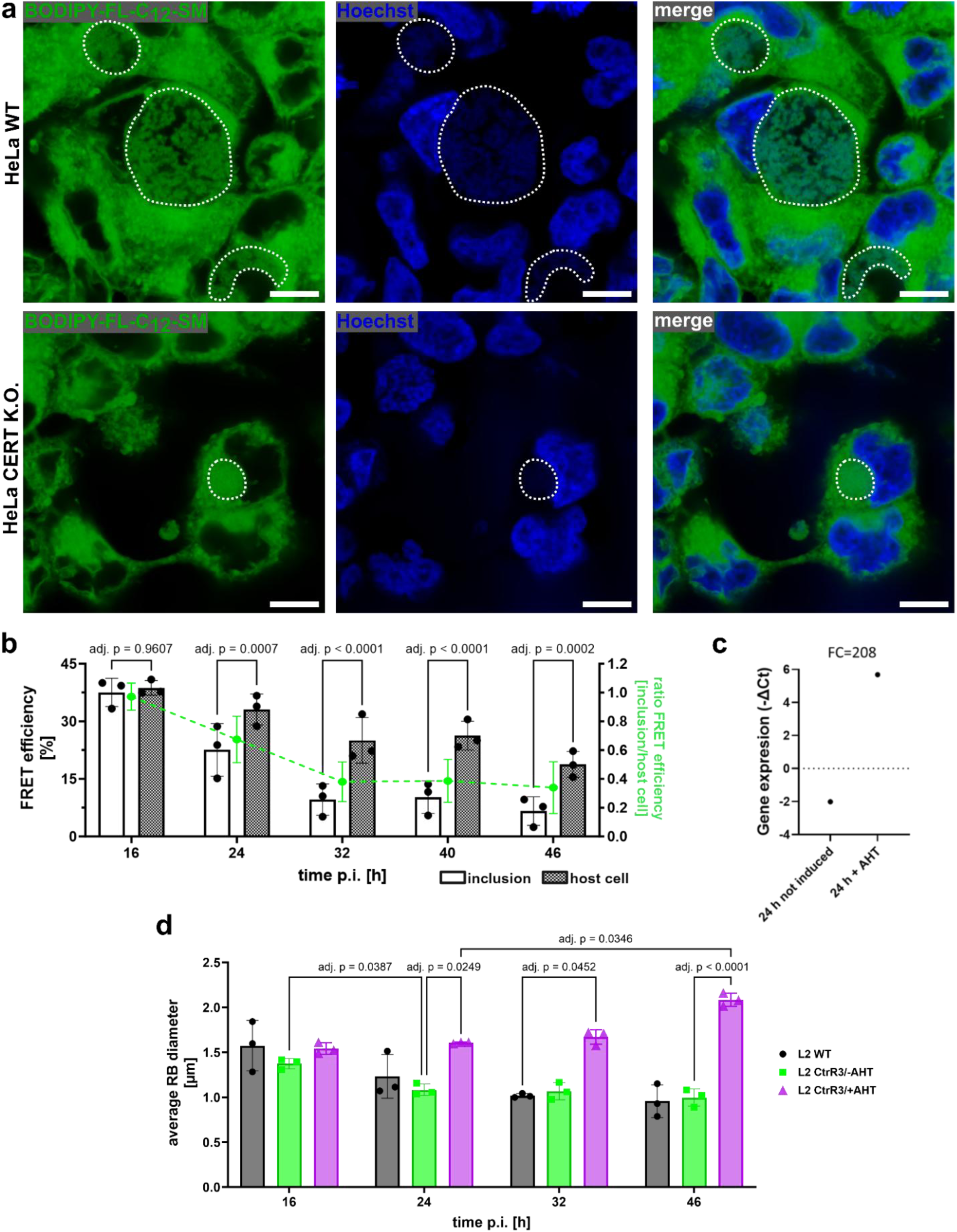
Measurement of TFSM metabolization in 4x ExM samples and characterization of the *C. trachomatis* L2 CtrR3 stain. **a, BODIPY-FL-C_12_-SM is taken up by inclusions.** HeLa cells were infected with *C. trachomatis*. After 3h, cells were treated with 10 µM BODIPY-FL-C_12_-SM and fixed 40 h p.i.. DNA was stained by Hoechst. Inclusions are indicated with white lines. Scale bar: 10 µm. **b, TFSM degradation in inclusions can be measured by 4x ExM.** HeLa 229 cells were infected with *C. trachomatis* in presence of TFSM2 for indicated time points and 4-fold expanded. FRET efficiency in inclusions and host areas were determined by FRET acceptor bleaching. FRET efficiency ratios (inclusions vs. host cell) were calculated (green). **c, CtrR3 is overexpressed under control of anhydrous tetracycline (AHT).** HeLa 229 cells were infected with the *C. trachomatis* CtrR3 strain in presence or absence of AHT for 24 h. CtrR3 expression was determined by RT-qPCR. **d, overexpression of CtrR3 affects the size of RBs.** HeLa 229 cells were infected with *C. trachomatis* L2 WT, or the CtrR3 strain either in absence or presence of AHT for the indicated time points. RBs were visualized by TFSM, and diameters were determined by 4x ExM. Statistics: Two-way ANOVA and Šidák’s multiple comparison (a) or Tukey’s multiple comparison (c).

**Supp. Fig. 6.**
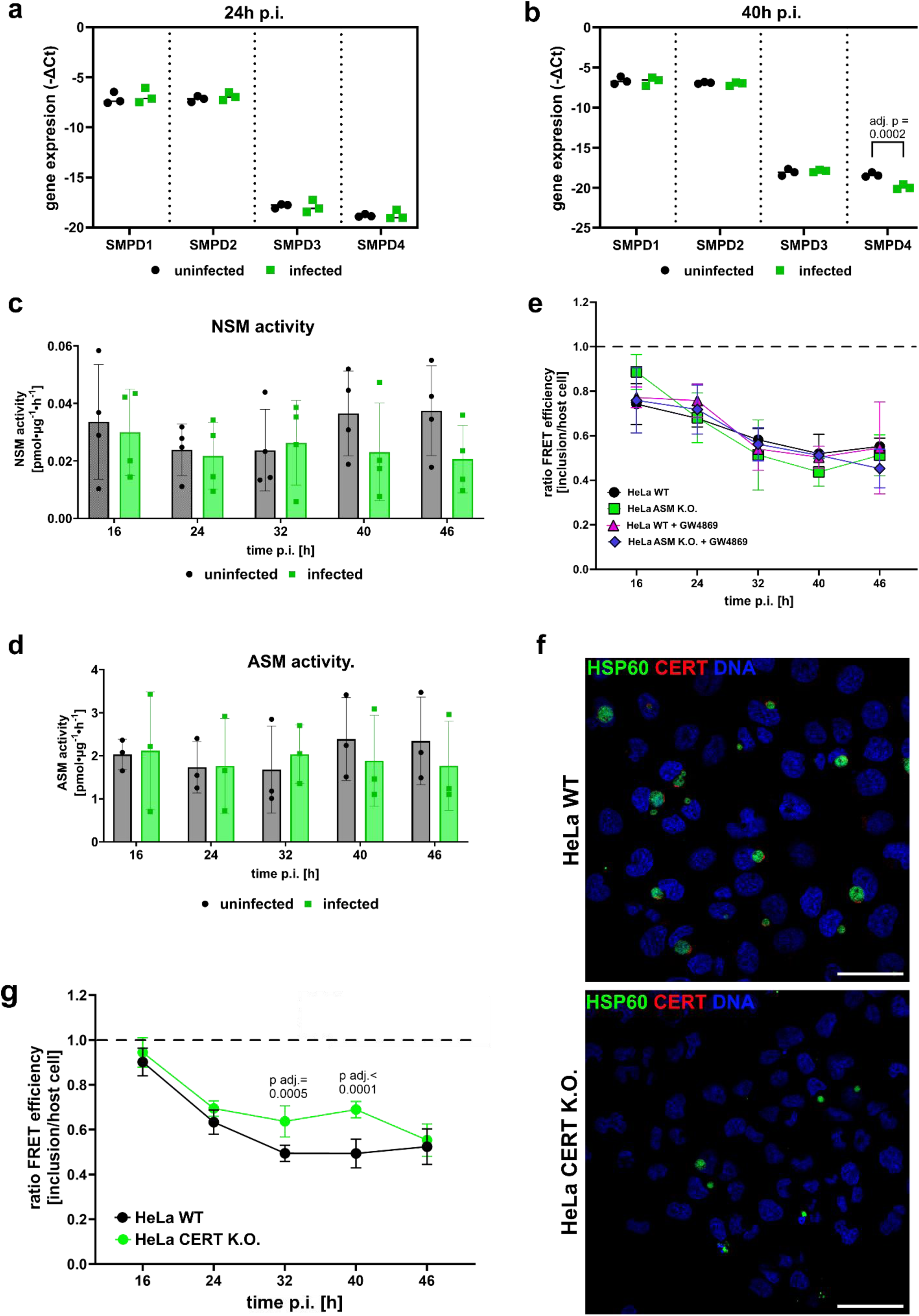
Accumulation of metabolized TFSM in inclusions is independent of major host sphingolipid metabolic factors. **a-d, host SMase expression levels and cellular SMase activity is not affected by *C. trachomatis* infection.** HeLa 229 WT cells were infected with *C. trachomatis*. At the indicated time point, either RNA was extracted and expression of host cell SMases (SMPD1-SMPD4) were measured by RT-qPCR (a, b) or cells were lysed and cellular neutral sphingomyelinase (NSM, c) as well as acid sphingomyelinase (ASM, d) activity was measured. n≥3. **e, accumulation of metabolized TFSM in inclusions is independent from host ASM and NSM.** HeLa WT or HeLa ASM K.O. were infected with *C. trachomatis* and if indicated treated with the NSM inhibitor GW4869 in presence of TFSM2. At the indicated time point, samples were fixed, TFSM was stained with BOD-IPY-FL-DBCO and AZDye546-azide, 4-fold expanded and FRET efficiencies in inclusions and host cell areas were determined. FRET efficiency ratios “inclusion vs. host cell” were calculated. n≥ 3. **e, absence of CERT impairs infection establishment but does not affect the accumulation of metabolized TFSM in inclusions.** HeLa WT or HeLa CERT K.O. cells were infected with *C. trachomatis* in presence of TFSM2. At the indicated time point, samples were fixed, and either immunostained for chlamydial HSP60, CERT and DNA (e) or stained for TFSM2, 4-fold expanded and FRET efficiency ratios “inclusion vs. host cell” were determined (g). Scale bar: 50 µm. n≥3. Statistics: Two-way RM ANOVA and Šidák’s multiple comparison.

**Supp. Fig. 7.**
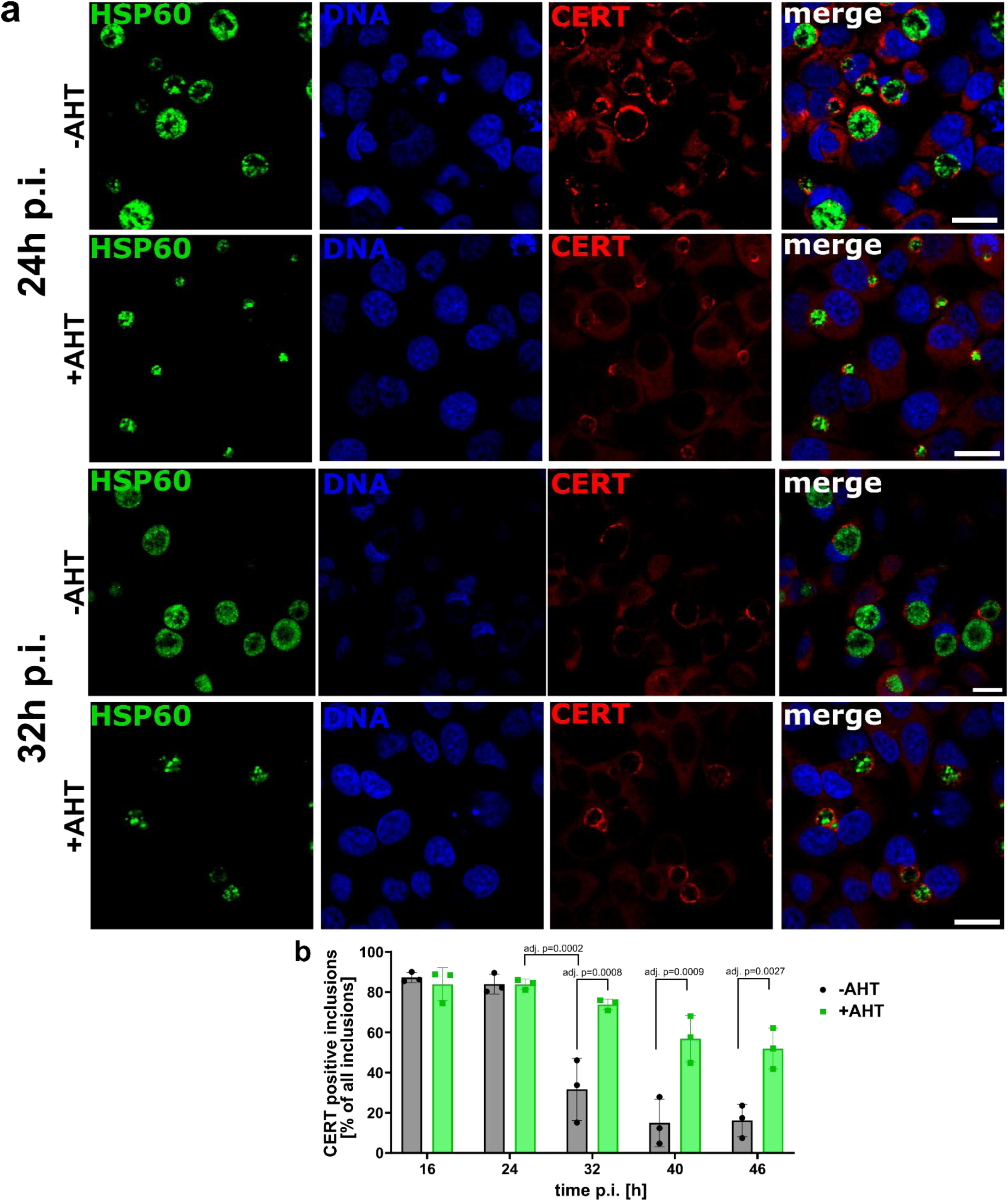
CERT is recruited to inclusions of *C. trachomatis* overexpressing the sRNA CtrR3. HeLa 229 were infected with *C. trachomatis* L2 WT or the L2 CtrR3 strain either in presence or absence of AHT for the indicated duration. Cells were fixed, stained with an anti-CERT (red) as well as an anti-chlamydial HSP60 (green) antibody and recorded by conventional confocal microscopy. The proportion of CERT-positive inclusions was determined (b). Scale bars: 20 µm. Statistics: Two-way RM ANOVA and Tukey’s multiple comparison.

**Supp. Fig. 8.**
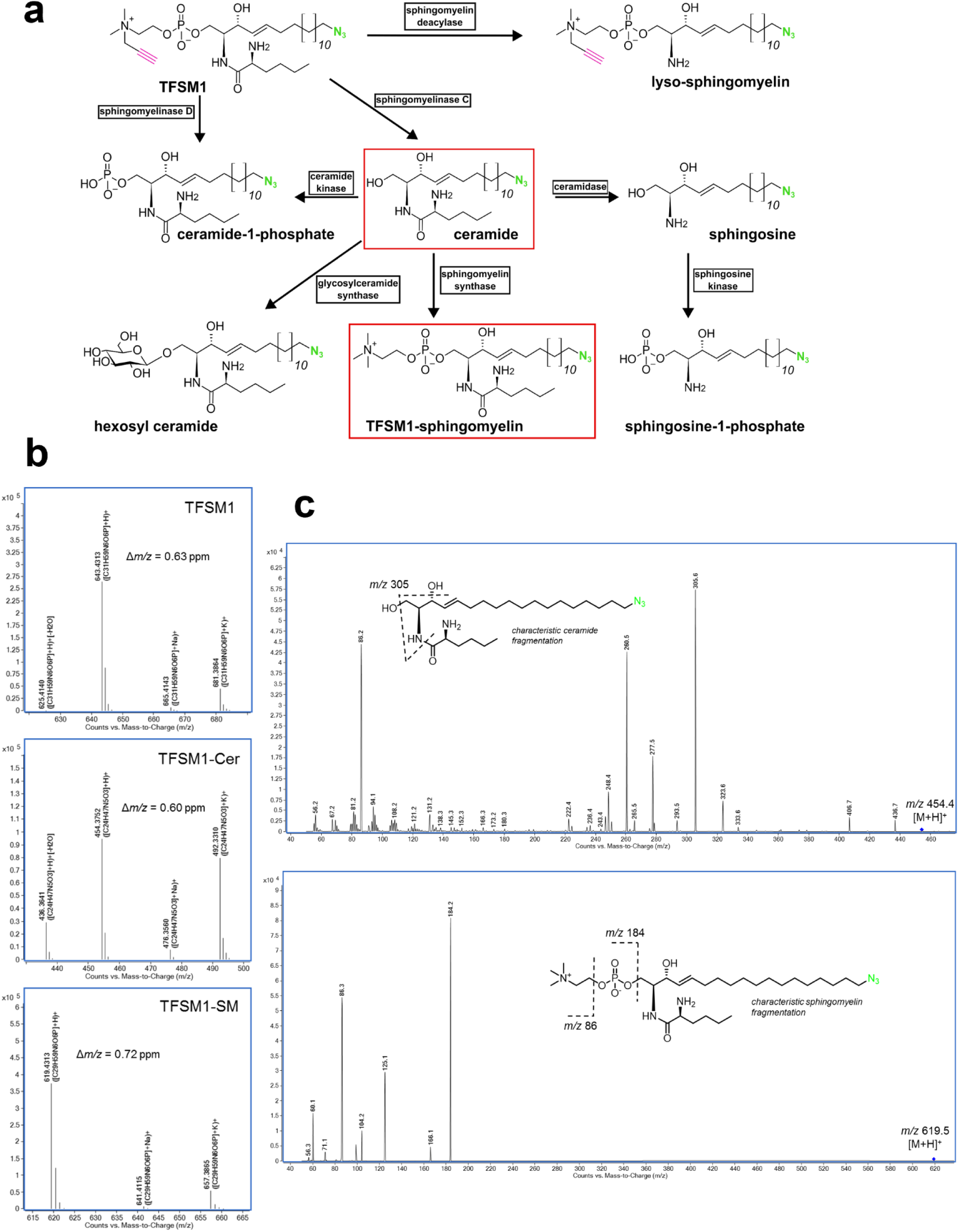
Screening for TFSM metabolites during *C. trachomatis* infection. (a) HeLa 229 cells were treated with TFSM1 and either infected with *C. trachomatis* in the presence of TFSM1 or left uninfected. After 24 h or 40 h p.i. lipids were extracted and analyzed for the indicated TFSM1 metabolites. Detected metabolites were highlighted in red. (b), Identification of the assay substrate TFSM1 and its metabolites TFSM1-Cer and TFSM1-SM based on the accurate masses (*m/z*) of molecular ions determined by high-resolution QTOF mass spectrometry. (c), Representative product ion (MS/MS) spectra of TFSM1-Cer (top, recorded with 20 eV collision energy) and TFSM1-SM (bottom, recorded with 60 eV collision energy), which exhibit characteristic MS/MS fragmentation patterns of ceramide and sphingomyelin species, respectively (illustrated with the help of the structural formula as insets).

## Supplementary Movies

**Supp. Movie 1 Z-stack of HeLa cells infected with *C. trachomatis* 1 h p.i. and visualized by 8x ExM.**HeLa 229 cells were preloaded with TFSM2, infected with *C. trachomatis* in presence of TFSM2, and fixed 1 h p.i.. Samples were stained with BOIDPY-FL-DBCO as well as AZDye546-azide and 8-fold expanded. Z stacks of BODIPY-FL (green) and FRET (red) channels were recorded on a Leica Stellaris 5 microscope. Scale bar: 16 µm (2 µm with 8x expansion factor).

**Supp. Movie 2 Cryo-tomogram followed by a segmentation of a cryo-FIB milled EB particle.**Green, membrane stack; blue, nucleoid; red, T3SSs; black, outer membrane, dark grey; inner membrane; light grey, cytoplasm. Scale bar: 200 nm.

## Supplementary Notes

**Supplementary Note 1**

TFSM1 and TFSM2 differ by the positioning of the azido function (**Supp. Fig. 1**). While the azido function in TFSM1 is located in sphingoid backbone, in TFSM2 it can be found in the acyl chain. Both molecules enable monitoring of SM metabolization by microscopy and lipidomics. TFSM1 is metabolized slower than TFSM2 in HEK cells.^45^ In general, both compounds behaved similar during *C. trachomatis* infection.^45, 46, 106^ However, dependent on the application the positioning of the azido function can be important.

TFSM1 is ideal for monitoring sphingolipid metabolization by lipidomics as the azido function in the sphingoid backbone can be used to detect the TFSM1-derived sphingosine metabolite when the acyl chain is cleaved off, e.g. by ceramidases (see **Supp. Fig. 8**). In contrast, TFSM2-derived sphingosine is indistinguishable from naturally occurring sphingosine.

Staining TFSM2 with DBCO dyes may be more efficient than TFSM1 as the azido function might be more accessible for the dye when positioned on the shorter acyl chain. Hence, we prefer using TFSM2 when monitoring sphingolipid distribution via 8x ExM.

In theory, FRET originating from TFSM2 can only be detected when the ceramide tail and the head group is present in the same molecule. By contrast, TFSM1 that was processed to lyso-sphingomyelin by a sphingomyelin deacylase can still result in a FRET signal (see **Supp. Fig. 8**).

**Supplementary Note 2**

To exclude that the accumulation is caused by anisotropic sample expansion, we compared EBs recorded by 4x and 8x ExM and stained with TFSM2 (**Supp. Fig. 2,** a, b). We determined the diameter of the accumulation and of whole EBs and calculated the ratio (**Supp. Fig. 2**, c). The relation between accumulation and EB diameter was similar in 4-fold and 8-fold expanded samples, indicating isotropic sample expansion and excluding expansion artifacts (**Supp. Fig. 2**, d). We also compared the diameter of the accumulation, whole EBs and RBs in 4x- and 8x-expanded samples (**Supp. Fig. 2**, e). In theory, it is expected that structures appear 2-fold larger in 8x ExM compared to 4x ExM images. However, the average accumulations diameter increased by ∼1.6-fold, EB diameter by ∼1.5-fold and RB diameters by ∼1.8-fold (**Supp. Fig. 2**, f). The slight differences in expansion of different structures are statistically not significant, indicating that the sample expands largely isotropically.

The click dyes that we use to label TFSM2 can introduce comparably high background staining. To measure the contribution of this background to the detected signal, we infected samples with *C. trachomatis* in presence or absence of TFSM2. Then, we stained the samples with varying concentrations of the click dye and performed 4x or 8x ExM. We compared the fluorescence intensity within the accumulation in samples incubated with or without TFSM2 and determined the signal-to-background ratio (**Supp. Fig. 2**, g). For dye concentrations commonly used in our expansion protocols, an average signal-to-background ratio of ∼3:1 was determined, demonstrating that the accumulation truly contains TFSM2.

**Supplementary Note 3**

For TFSM2 staining, HeLa 229 cells were infected with *C. trachomatis* in presence or absence of TFSM2 and samples were fixed at 32 h p.i..

For staining with α-NH_2_-ω-N_3_-C_6_-ceramide, cells were infected with *C. trachomatis* for 31 h and then α-NH_2_-ω-N_3_-C_6_-ceramide was added for 1 h before samples were fixed.

Functionalized lipids were stained with BODIPY-FL-DBCO and samples were expanded 8-fold. DNA and proteins were visualized by Hoechst and NHS ester staining. For measuring lipid content in chlamydial nucleoids, areas containing nucleoids were selected based on Hoechst signal and the signal intensities in all three channels were measured. Mean fluorescence intensities measured in individual nucleoids were plotted (see **Supp. Fig. 3**, a for one representative replicate) and the average fluorescence intensities of several independent biological replicates were determined. Ratios between results obtained from samples incubated with the lipid derivatives and samples without added analogs (but stained with the DBCO dye) were determined (**Supp. Fig. 3**, b). We detected a significantly higher BODIPY-FL signal in nucleoids recorded in samples incubated with the functionalized lipids compared to staining controls.

TFSM2 is taken up by host cell via endocytosis, due to its polar head group. By contrast, α-NH_2_-ω-N_3_-C_6_-ceramide is membrane permeable and thus, it is rapidly incorporated into cellular membranes. Hence, shorter incubation times are required for α-NH_2_-ω-N_3_-C_6_-ceramide (1h) compared to TFSM (several hours).

**Supplementary Note 4**

RBs of WT and the uninduced *C. trachomatis* L2 CtrR3 strain possessed the highest diameter at 16 h p.i. (∼1.5 µm) and shrunk during infection to an average size of ∼1 µm (**Supp. Fig. 5**, d). By contrast, the diameter of RBs upon CtrR3 overexpression remained at their initial size up to 32 h p.i. and even increased to ∼2 µm at 40 h p.i., suggesting a connection between RB-EB redifferentiation and RB size as described previously.^34^

**Supplementary Note 5**

Since *C. trachomatis* often modulates host cell metabolism for adequate nutrient supply,^138^ we hypothesized that TFSM might be metabolized by host cell enzymes and is then shuttled into the inclusion. Human cells possess three neutral (NSM1-3)^139^ and one acid SMases (ASM)^16^, which catalyze the metabolization of SM to ceramide. We neither detected increased expression levels of these enzymes (**Supp. Fig. 6**, a and b), nor did we measure an increased cellular NSM (**Supp. Fig. 6**, c), or ASM (**Supp. Fig. 6**, d) activity upon infection, suggesting that host cell SMases are negligible for the enrichment of metabolized TFSMs in chlamydial inclusions.

The ceramide transfer protein (CERT) was previously shown to be recruited to chlamydial inclusions^108^, which could cause the enrichment of metabolized TFSM within inclusion. To this end, we infected a HeLa CERT K.O. cell line^109^ with *C. trachomatis* and again performed FRET acceptor bleaching to determine native TFSM content in inclusions and host cell membranes. Infection rates were strikingly lower in CERT K.O.s compared to the wild type cells (**Supp. Fig. 6**, e). However, metabolized TFSMs accumulated in inclusions despite the absence of CERT, even though in an attenuated fashion (**Supp. Fig. 6**, f).

We also compared recruitment of CERT to inclusions in cells infected with *C. trachomatis* L2 CtrR3 in presence or absence of AHT. Consistently, CtrR3 expression, which strongly attenuated the enrichment of metabolized TFSMs in inclusions (**Fig. 5**, e), did not interfere with the recruitment of CERT (**Supp. Fig. 7**, a and b). A large proportion of inclusions were positive for CERT at 16 and 24 h p.i. independently from CtrR3 expression. Interestingly, the proportion of CERT-positive inclusions decreased in samples without CtrR3 expression at 32h p.i., whereas most inclusions remained CERT-positive if CtrR3 was expressed.

In summary, our data indicates that TFSM metabolization during *C. trachomatis* infection is independent from central factors of host cell sphingolipid metabolism.

